# Endogenous lipid droplets exhibit profound proteomic and lipidomic differences across neural stem cell states

**DOI:** 10.64898/2026.08.18.745393

**Authors:** Diana Panfilova, Mergim Ramosaj, Manfredo Quadroni, Marlen Knobloch

**Affiliations:** Department of Biomedical Sciences, University of Lausanne, Lausanne, Switzerland; Protein Analysis Facility, University of Lausanne, Lausanne, Switzerland

## Abstract

Lipid droplets (LDs) are protein-coated organelles that store neutral lipids and regulate diverse cellular processes beyond energy metabolism. In neural stem/progenitor cells (NSPCs), LD abundance and morphology vary across cellular states, yet whether LD molecular composition is similarly state-dependent remains unknown. Here, we define the first endogenous LD proteome and lipidome atlas of NSPCs and their progeny. State-resolved analyses reveal extensive differences in both LD-associated proteins and stored lipids, allowing for identification of LD signatures that distinguish quiescent and proliferative states, and uncovering selective enrichment of numerous proteins on quiescent NSPC LDs. Functional interrogation of one such protein, CIDEB, showed that its knockdown alters LD morphology and induces senescence-associated transcriptional programs, implicating CIDEB in the maintenance of NSPC quiescence. These findings establish LDs as dynamically specialized organelles in NSPCs and their progeny and provide a resource for investigating LD-mediated regulation of stem cell state and lineage progression.

## Introduction

Adult neurogenesis is the process of generating new neurons throughout life and is linked to cognition and memory formation in mammals (*1*). This process is restricted to distinct brain areas, called neurogenic niches, with the subventricular zone (SVZ) and the hippocampal dentate gyrus (DG) being the main sites of postnatal neurogenesis in rodents (*2*), and ample evidence for neurogenesis in the adult human DG (*3–5*). Newborn neurons arise from neural stem/progenitor cells (NSPCs) that reside in the neurogenic niches in a quiescent state (qNSPCs). qNSPCs can be activated by certain stimuli, gaining capacity to proliferate (prolNSPCs, also known as activated NSPCs) and subsequently differentiate into neurons or glial cells such as astrocytes or oligodendrocytes (*6*). NSPC state and activity are tightly regulated by metabolic processes, and accumulating evidence shows that lipid metabolism plays a crucial role in NSPC regulation (*7*, *8*). In particular, qNSPCs have higher fatty acid beta-oxidation (FAO) levels compared to prolNSPCs, which depend on *de novo* lipogenesis, showing that shifting the balance between lipid build-up and breakdown can regulate NSPC states (*7*, *8*).

Lipid droplets (LDs), long perceived as inert cytoplasmic fat inclusions, have been established in recent years as complex organelles with key functions in lipid homeostasis. LDs store neutral lipids, mainly triacylglycerides (TGs) and cholesterol esters (CEs), in a core surrounded by a phospholipid monolayer. The LD surface is coated by so-called LD-coat proteins that can regulate LD stability and turnover (*9*). Classical examples include perilipin 1-5 (PLIN1-5) family members that, among other functions, regulate LD stability and turnover (*10*). Another LD protein family, cell death-inducing DFFA-like effectors A-C (CIDE A-C), facilitate fusion of free-floating LDs, supporting their growth in adipose tissue (CIDEA and C) and hepatocytes (CIDEB) (*11*).

LDs occur in nearly all cell types. Depending on metabolic condition and health, LDs have been detected in adipose tissue, liver, skeletal muscle, retina, and brain (*12–16*). In the brain, LDs were long believed to form only under disease-related conditions such as neurodegeneration and aging (*17*). However, our recent studies show that LDs also accumulate in various cell types of the young healthy mouse brain, including NSPCs (*18*, *19*).

We previously showed that LD availability affects prolNSPC metabolism and proliferation, and that LD morphology varies by NSPC state: prolNSPCs have uniformly small LDs, whereas qNSPCs form a subset of large-diameter LDs (*9*). However, whether these morphological differences relate to LD function in NSPCs or reflect differences in LD lipid core and LD-coat protein composition across NSPC states remains unclear.

LD-coat proteins are key determinants of LD functions in a cell and contribute to roles far beyond lipid storage. For example, in *Drosophila* embryos LDs store and sequester excess histones to prevent their degradation and protect cells from a histone overload (*20–22*). LD-core lipid composition can also reflect LD functions: LDs sequester polyunsaturated fatty acids in form of TGs in their core to protect them from peroxidation and prevent fatty acid-induced lipotoxicity such as ferroptosis (*23*). Thus, during the last decade the scientific community has begun establishing LD proteomic and lipidomic datasets to better understand LD functions across cell types (*24–26*). However, generating such datasets is challenging for a few reasons: 1) LDs form membrane contact sites (MCSs) with multiple organelles (*27*), causing coextraction during LD isolation and complicating *bona fide* LD proteome identification; 2) many cultured cell types form LDs only upon fatty acid loading, limiting the creation of endogenous LD datasets. Moreover, LD structure has been shown to vary by cell type (*24*, *26*), underscoring the need for structural datasets across diverse cells. To date, the LD research community has created a web resource (LD knowledge portal, www.lipiddroplet.org) listing *bona fide* LD proteins identified using advanced methods to obtain high quality LD proteome (i.e., APEX2-proximity labeling (*24*), protein correlation profiling (*25*), comparison with total cell proteome (*26*)) in four human cell lines and mouse liver. However, no studies have profiled endogenous LDs in stem cells.

In this study we isolated endogenous LDs from adult murine qNSPCs, prolNSPCs, and NSPC-derived astrocytes, and established their proteome and lipidome to further understand the role of LDs in these cells. We found that both LD proteomes and lipidomes were highly cell-type specific, with many proteins specifically enriched on qNSPC LDs. Among these qNSPC-specific proteins, we identified CIDEA and CIDEB and showed that CIDEB depletion altered LD size and induced a senescence-like phenotype in qNSPCs.

## Results

### Establishing a high-confidence LD proteome of endogenous LDs from different NSPC states

Primary adult NSPCs in culture can be kept under quiescence conditions (qNSPCs), proliferative conditions (prolNSPCs), or allowed to differentiate into astrocytes and neurons through growth factor withdrawal (diffNSPCs), recapitulating the main NSPC states present during adult neurogenesis (Fig. 1A). Characterization of LDs in NSPCs derived from the SVZ of tdTom-Plin2 mice, an endogenous LD-reporter mouse line that allows for LD visualization without staining procedure (*18*), showed clear differences in LD morphology between these cell types (Fig. 1B). While we did not observe statistically significant changes in average LD volume, total LD volume, or LD numbers, diffNSPCs tended to have more LDs and bigger total LD volume compared to qNSPCs and prolNSPCs (fig. S1A-C). However, qNSPCs formed a subset of LDs that were significantly larger than those in prolNSPCs or diffNSPCs (Fig. 1C), in line with our previous results obtained using PLIN2 staining in NSPCs derived from wild-type (WT) mice (*9*).

**Fig. 1.**
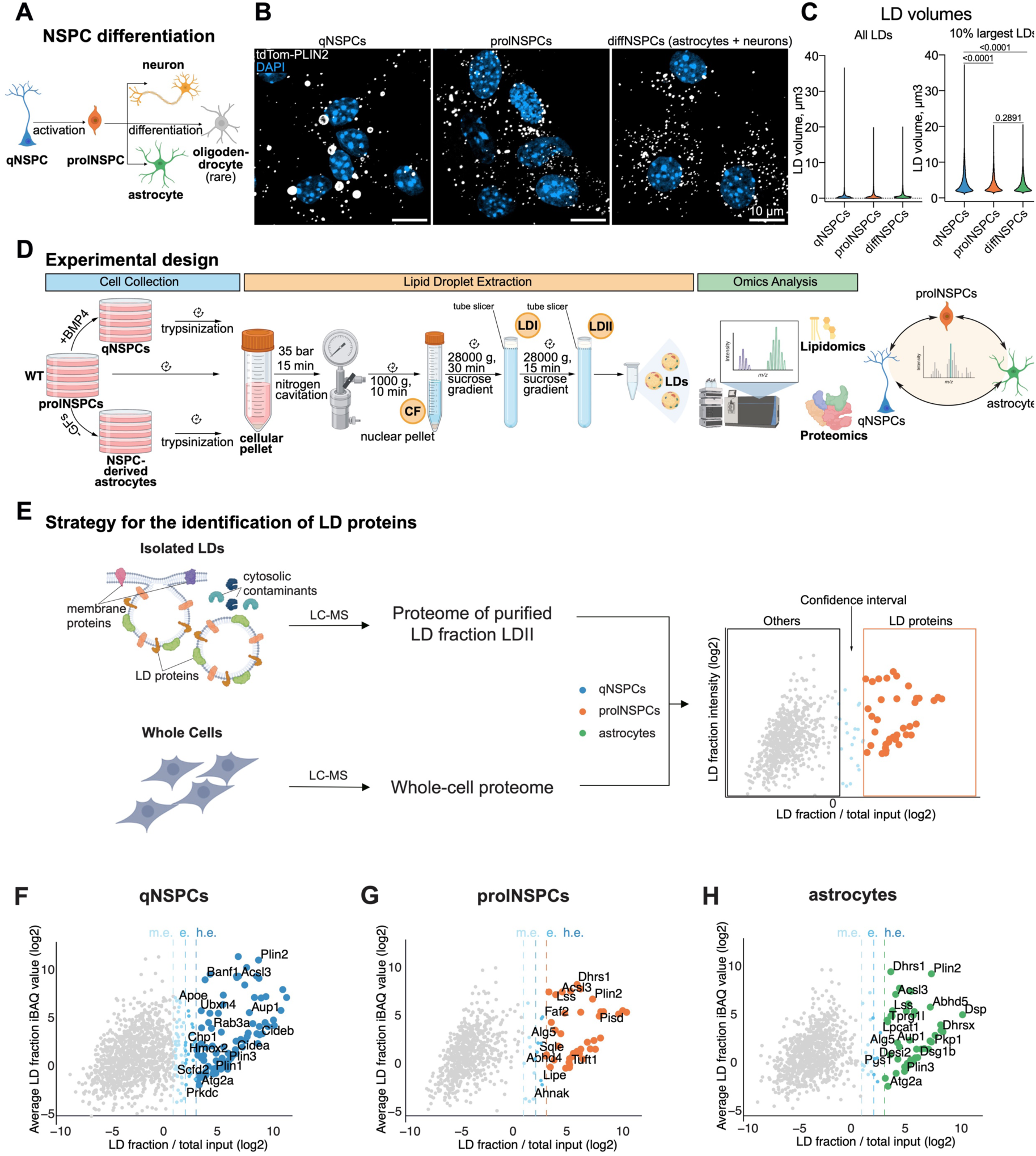
LD extraction from NSPCs and their progeny and identification of high-confidence LD proteins. (A) A simplified scheme of neural stem/progenitor cell (NSPC) differentiation during adult neurogenesis. qNSPC: quiescent NSPC, prolNSPC: proliferative NSPCs. (B) Representative confocal images of qNSPCs, prolNSPCs and differentiating NSPCs (diffNSPCs: co-culture of astrocytes and neurons), derived from the subventricular zone of 8-week-old tdTom-Plin2+/- mice, show lipid droplet (LD) morphology of the cell types (tdTom-PLIN2: LDs, DAPI: nuclei). Representative images are maximum-intensity projections. (C) Violin plot showing LD-volume distribution of LDs of all sizes (left) and top 10% of largest LDs (right) in qNSPCs, prolNSPCs, and diffNSPCs. Data collected from three independent experiments. Anderson-Darling k-sample test with Holm correction. (D) Experimental design for LD purification from wild-type (WT) qNSPCs, prolNSPCs, and NSPC-derived astrocytes for liquid chromatography-coupled mass-spectrometry (LC-MS) analysis. GFs: growth factors, BMP4: bone morphogenetic protein 4, CF: cytosolic fraction, LDI: LD fraction I, LDII: LD fraction II. (E) Strategy for identifying high-confidence LD proteomes. Alongside LD fractions, whole-cell lysates were isolated in a separate experiment to analyze total-cell proteomes by LC-MS. Proteins were defined as LD-resident if log2FC of protein abundance in LD fraction relative to total input exceeded the cut-off threshold (orange box). Confidence interval is defined by the cut-off and comprises the proteins that fall between non-enriched proteins and specifically enriched proteins. (F-H) Scatter plots showing log2FC of protein abundance in the LD fraction versus whole-cell proteome and average LD-fraction abundance (intensity-based absolute quantification, iBAQ). Proteins with LD-fraction/total-input log2FC < 1 are gray, moderately enriched (m.e.) proteins with 1 ≤ log2FC < 2 are light blue, enriched (e.) proteins with 2 ≤ log2FC < 3 are blue and highly enriched (h.e.) proteins with log2FC ≥ 3 are dark blue (qNSPCs, F), orange (prolNSPCs, G), or green (NSPC-derived astrocytes, H). FC: fold change. Protein quantification is based on four biological replicates.

We next asked whether these differences in LD morphology across NSPC states could reflect differences in LD protein and lipid composition. Therefore, we adapted an LD-isolation protocol using nitrogen cavitation and density-gradient centrifugation to obtain a pure LD fraction for proteomic and lipidomic analysis (Fig. 1D) (*28*). We used primary WT SVZ NSPCs cultured as qNSPCs, prolNSPCs and diffNSPCs. To obtain a homogeneous diffNSPC population, we washed off the neurons prior to LD collection, as we previously showed that neurons have far fewer LDs than astrocytes (*9*).

For four independent experiments per cell type, we lysed cellular pellets coming from approximately 100 million cells using nitrogen cavitation and used the cytosolic fraction (CF) for ultracentrifugation in a sucrose density gradient. We collected the first LD fraction (LDI) and repeated the ultracentrifugation to further purify LDs from cytosolic and membrane contaminants. After the second round of density-gradient step, we collected the purified LD fraction (LDII).

We validated LD-fraction purity using the neutral-lipid dye Bodipy493/503 and Western blotting against PLIN2 (fig. S1D, E). While both fractions contained LDs, the LDII fraction showed absence of cytosolic protein beta-ACTIN signal, indicating increased purity relative to LDI (fig. S1E). We therefore analyzed the proteome and lipidome of LDII fractions from qNSPCs, prolNSPCs and NSPC-derived astrocytes using liquid chromatography-coupled tandem mass spectrometry (LC-MS/MS) with label-free quantification for LD proteins and hydrophilic interaction liquid chromatography coupled to electrospray ionization tandem mass spectrometry (HILIC-ESI-MS/MS) for LD lipids.

LDs are highly interconnected with other organelles, making purification challenging. Although our LDII fraction showed high purity level on Western blot, additional steps were needed to establish a high-confidence LD proteome and rule out non-LD proteins that might co-purify.

Indeed, in LDII fraction we identified 1425 proteins by LC-MS analysis, whereas usual *bona fide* LD proteomes contain ∼100 proteins (table S1). To filter out potential non-LD proteins, we used the strategy described by Mejhert et al. to narrow down LD-enriched proteins (Fig. 1E) (*26*). We analyzed whole-cell lysate proteomes of qNSPCs, prolNSPCs and NSPC-derived astrocytes (table S1) and filtered out proteins not specifically enriched in the corresponding LD fraction (Fig. 1F-H), using three different fold-change (FC) cut-offs: moderately enriched (m.e.), with 2-4 fold increase (1 ≤ log2FC <2); enriched (e.), with 4-8 fold increase (2 ≤ log2FC <3 LD); and highly enriched (h.e.), with ≥ 8 fold increase (log2FC ≥ 3) (table S2).

Using the criteria for high enrichment (≥ 8 fold increase), we found 86 proteins to be highly enriched on LDs of qNSPCs, 40 proteins on LDs of prolNSPCs, and 43 proteins on LDs of NSPC-derived astrocytes.

### High-confidence endogenous LD proteome shows cell-state specific differences in LD proteins, especially in qNSPCs

Previous studies have identified LD proteomes in various cell types after oleic-acid loading and in mouse liver in different metabolic conditions, and these results have been compiled into the LD knowledge portal (www.lipiddroplet.org) (*29*). Using this database, we created a reference list of LD proteins that included 310 *bona fide* proteins present in at least one of these LD-proteome datasets (table S3). Comparison of this reference list with our NSPC LD proteomes showed that at least 50% of the highly enriched LD proteins in our cells were already annotated as LD proteins on the portal (Fig. 2A). This shows that our endogenous LD-enrichment approach effectively detects classical LD proteins, including PLIN1-3, long-chain-fatty-acid-CoA ligases 3 and 4 (ACSL3-4), hormone-sensitive lipase (LIPE), and lanosterol synthase (LSS) (table S2).

**Fig. 2.**
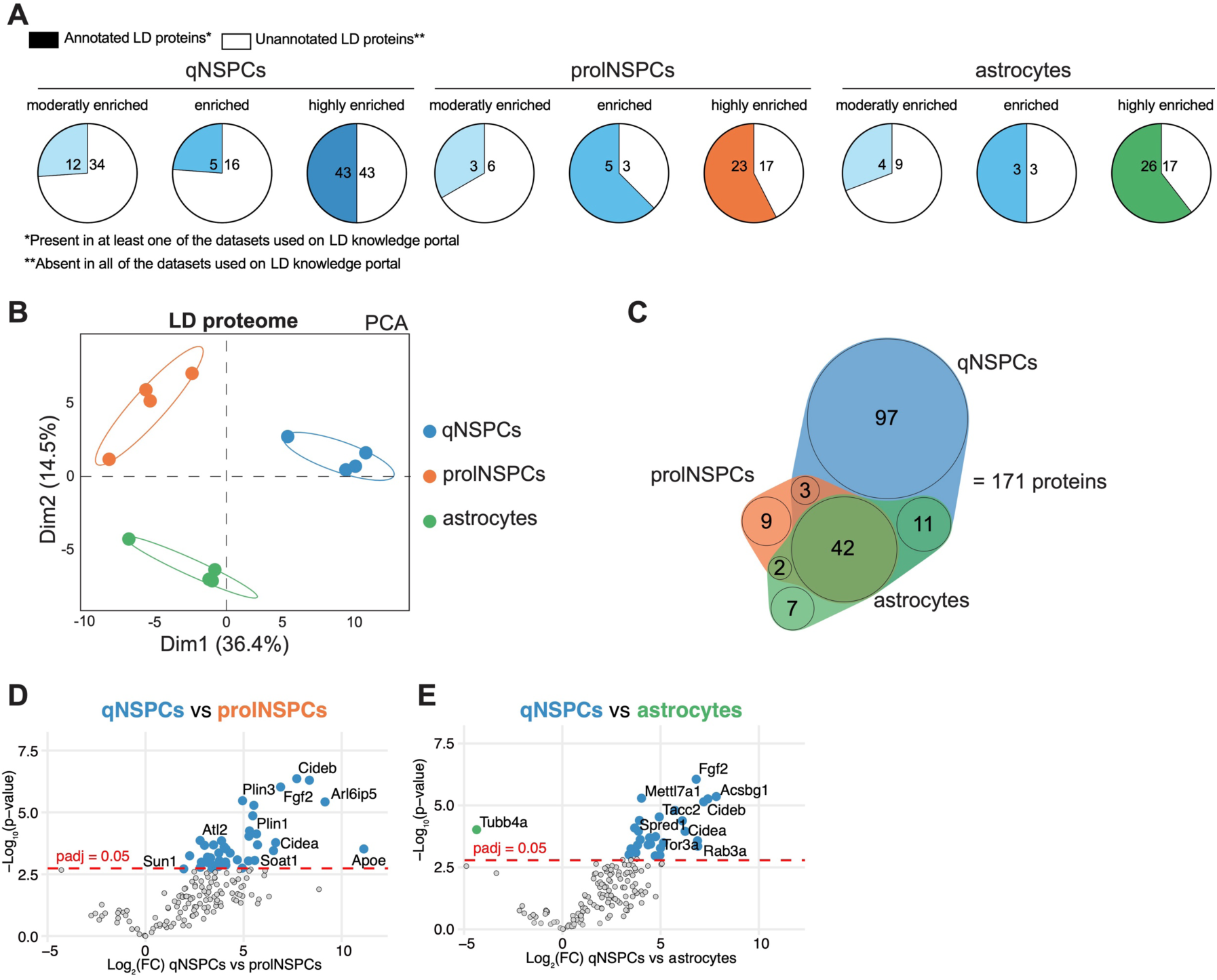
Cell-type-specific differences in the LD proteomes of qNSPCs, prolNSPCs and NSPC-derived astrocytes. (A) Pie-chart diagrams showing the number of proteins previously annotated to localize on lipid droplets (LDs) of neural stem/progenitor cells (NSPCs) in moderately enriched (1 ≤ log2FC < 2, light blue), enriched (2 ≤ log2FC < 3, blue) and highly enriched (log2FC ≥ 3, dark blue (qNSPC, left), orange (prolNSPC, middle), or green (NSPC-derived astrocytes, right)) categories, and those that have not been previously annotated (white). qNSPCs: quiescent s, prolNSPCs: proliferative NSPCs, FC: fold change. (B) Principal component analysis (PCA) of high-confidence LD proteins (log2FC ≥ 1 in at least one of the cell types) based on their abundance (C) Venn diagram of high-confidence LD proteins (log2FC ≥ 1 in at least one of the cell types). (D, E) Volcano plots based on high-confidence LD-protein quantification, depicting enrichment of LD proteins in qNSPCs compared to prolNSPCs (D) and in qNSPCs compared to NSPC-derived astrocytes (E). Significantly upregulated proteins are highlighted in dark blue for qNSPCs and green for NSPC-derived astrocytes. Gray dots represent proteins without significant abundance changes. The p-adjusted (padj) cut-off is 0.05 (multiple-comparisons adjusted). Protein quantification is based on four biological replicates.

Interestingly, many moderately enriched and enriched LD proteins were also previously annotated as LD proteins, including Ras-related (RAB) family proteins, acetyl-CoA carboxylase 1 (ACACA), and protein-L-histidine N-pros-methyltransferase (METLL9) (Fig. 2A, table S2). We speculate that some of these proteins may localize not only to LDs but also to other organelles or participate in LD MCS formation, resulting in lower log2FC values.

To further characterize LD-proteome differences between our cell types, we pooled all the proteins that were at least 2-fold enriched in the LD fraction in at least one cell type, yielding 171 LD proteins (fig. S2A, table S4) (we chose a 2-fold cut-off as we could see many previously annotated LD proteins to fall into this category). These proteins clearly separated qNSPCs, prolNSPCs, and NSPC-derived astrocytes in a principal component analysis (PCA), illustrating cell-state-specific LD-protein differences (Fig. 2B). Comparing these 171 proteins, we found 42 proteins present in all three cell types, 9 proteins specific to prolNSPCs (meaning that they were not detected or did not surpass filtering in the other two cell types), and 7 proteins specific to astrocytes (Fig. 2C, fig. S2A). Surprisingly, 97 proteins, which represent more than half of the LD-protein list, were qNSPC-specific (Fig. 2C, fig. S2A).

Importantly, protein assignment as cell-type-specific in the Venn diagram was based solely on whether a protein fulfilled our criteria for classification as an LD protein in a given cell type. To capture the quantitative differences, we performed pairwise differential abundance analyses of LD proteins between qNSPCs, prolNSPCs, and astrocytes (Fig. 2D, E, fig. S2B). This pairwise comparison allowed us to further narrow down significantly changed LD proteins, using an adjusted p-value of 0.05 cut-off. As expected, many LD proteins were significantly upregulated on qNSPC LDs in both its comparisons.

Together, these data show that LDs differ in composition depending on cell state, with differences especially pronounced in qNSPCs.

### Identified NSPC LD proteins exhibit broad functional diversity

To better understand the functional role of LD proteins in NSPCs and their progeny, we plotted the 171 high-confidence LD proteins according to their main functional role into different categories based on their UNIPROT annotation, reflecting different cellular functions, and indicated in which cell type they were present (Fig. 3). Most identified proteins fell into categories already associated with LDs: metabolism, vesicular trafficking, ubiquitin-related proteins, and others (*24*). However, some of the 171 proteins did not fit into these categories: their main functions were instead related to gene-expression regulation, immune response, and interestingly also proliferation and cell cycle. The latter included key regulators of cell cycle such as fibroblast growth factor 2 (FGF2), pleiotrophin (PTN) and F-box only protein 50 (NCCRP-1) (Fig. 3). These data suggest that LDs in different NSPC states harbor non-canonical proteins.

**Fig. 3.**
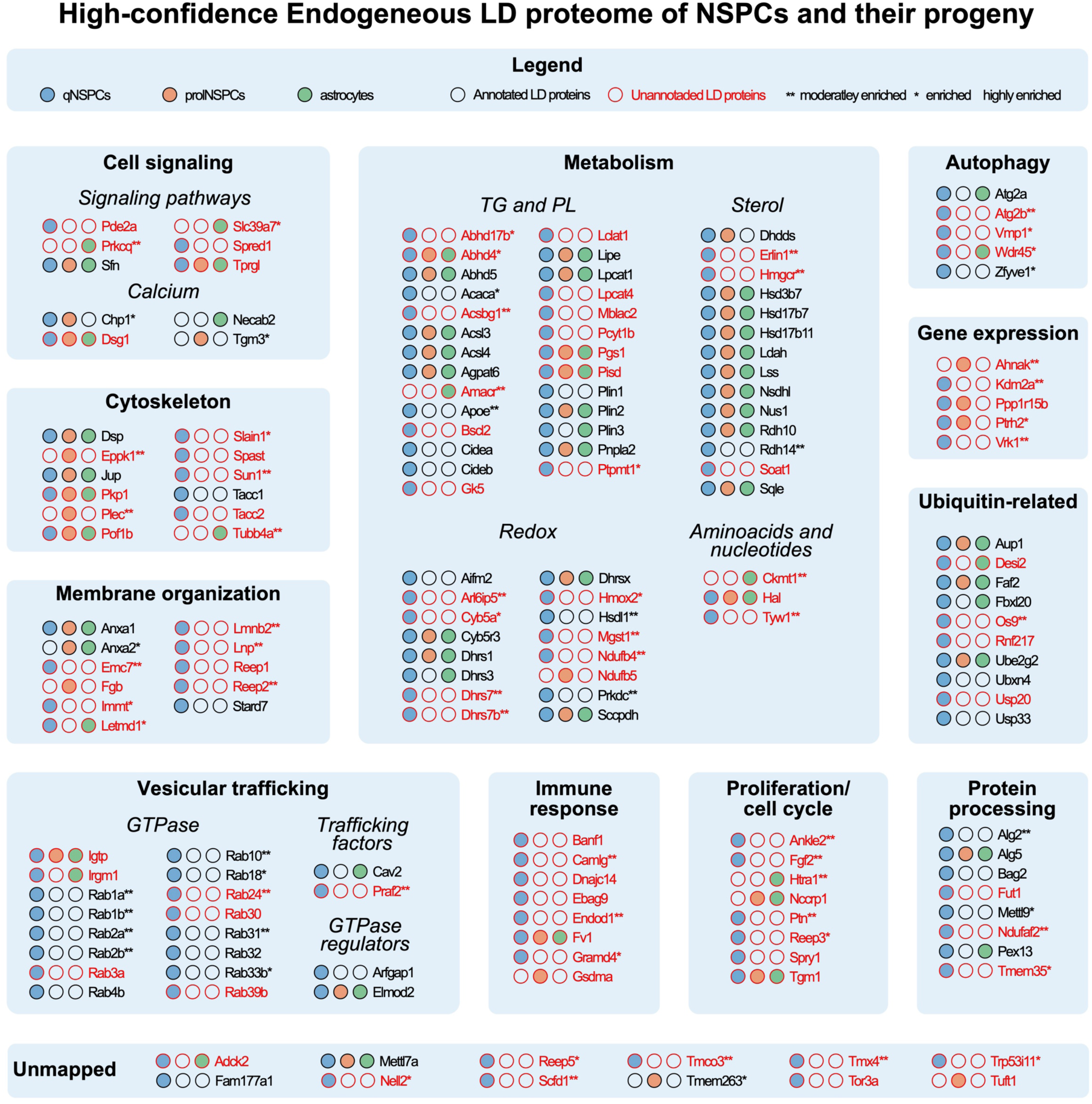
Combined high-confidence endogenous LD proteome of NSPCs and their progeny. Composite illustration of high-confidence endogenous lipid droplet (LD) proteins identified in quiescent neural stem/progenitor cells (qNSPCs, blue), proliferative NSPCs (prolNSPCs, orange) and NSPC-derived astrocytes (green). Proteins are grouped into functional modules based on UNIPROT annotation and color-coded red if they have not been identified as LD proteins in at least one dataset included in the LD knowledge portal. Asterisks indicate enrichment categories: ** for moderately enriched proteins (1 ≤ log2FC< 2), * for enriched proteins (2 ≤ log2FC < 3) and no asterisk for highly enriched proteins (3 ≤ log2FC). FC: fold change, TG: triacylglycerides, PL: phospholipids.

As expected, the majority of LD proteins in each functional category were detected in qNSPCs, with many proteins exclusive to qNSPCs (Fig. 3). For instance, all RAB-family proteins, which had been consistently detected across datasets in the LD knowledge portal, were localized only to LDs in qNSPCs. Interestingly, many identified LD proteins are also involved in establishing MCSs between LDs and other organelles (fig. S3).

To further explore potential differences in MCS proteins, we next assessed how many of the 171 proteins have been previously identified as LD inter-organelle regulators (*27*, *30–34*). We found 31 such proteins in our NSPC LD proteome: 11 proteins mentioned in multiple studies and 20 proteins recently identified by Guo and colleagues (*34*). We grouped these proteins into five categories based on the organelle-LD contact they were associated with: mitochondria-LD, LD-LD, endoplasmic reticulum (ER)-LD, lysosome-LD, and peroxisome-LD (fig. S3). While each of these groups, except for the peroxisome-LD group, contained proteins across all cell types, most were specific to qNSPCs, suggesting that qNSPCs may interact differently with other organelles than prolNSPCs and astrocytes. Importantly, because our dataset does not include information on the protein localization to other organelles, this analysis only provides a list of potential candidates for further investigations.

Summarizing, we established a novel dataset of high-confidence endogenous LD proteins in NSPCs and their progeny. This dataset highlights strong cell-type specificity in LD proteomic composition and provides a new resource to study LD-protein function in NSPCs.

### Endogenous LD lipidome also differs between cell states

Having established that the LD proteome was cell-type specific, we next analyzed whether the lipid content of the isolated LDs also differed between qNSPCs, prolNSPCs, and NSPC-derived astrocytes using targeted lipidomics. We identified 224 TG species, four CE species, five phosphatidylcholine (PC) species, six phosphatidylethanolamine (PE) species, and one lysophosphatidylcholin (LPC) species (table S5). Similar to LD proteins, these LD lipids clearly separated by the cell type in the PCA plot, showing that the LD lipidome is also cell-type specific (Fig. 4A). Interestingly, this separation was mostly explained by LD-core lipids TGs and CEs (Fig. 4B), while lipids from the LD phospholipid monolayer only separated prolNSPCs from qNSPCs and astrocytes (fig. S4A).

**Fig. 4.**
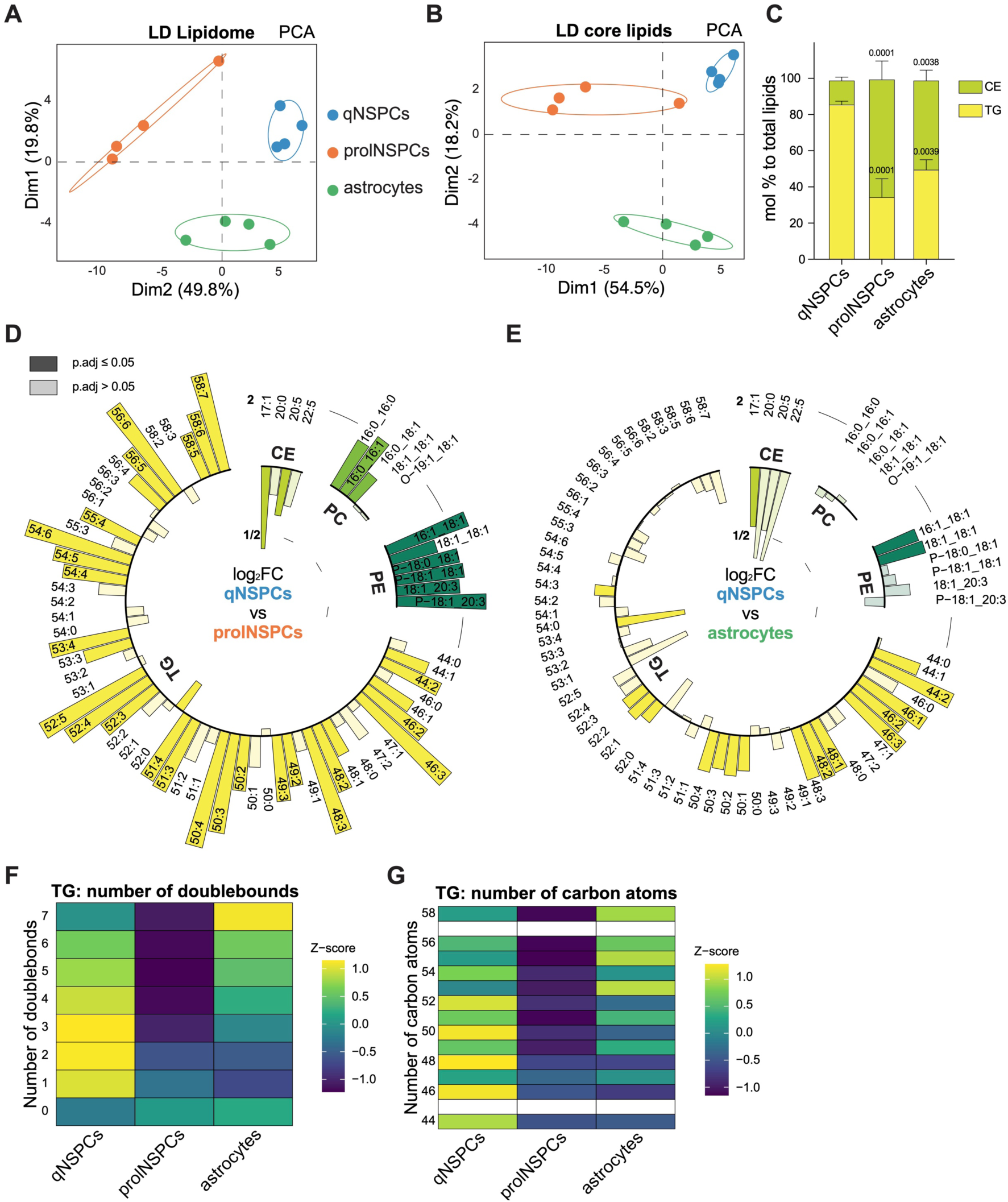
Cell-type specific differences in the LD lipidomes of qNSPCs, prolNSPCs and NSPC-derived astrocytes. (A, B) Principal component analysis (PCA) of lipid droplet (LD) samples based on the abundance of all identified lipids (A) and on the abundance of LD-core lipids triacylglycerides (TG) and cholesterol esters (CE) (B) in quiescent neural stem/progenitor cells (qNSPCs), proliferative NSPCs (prolNSPCs) and NSPC-derived astrocytes. (C) Distribution of LD-core lipid species detected in LDs of qNSPCs, prolNSPCs, and NSPC-derived astrocytes, color-coded by lipid class. %mol of total lipids (all detected lipid species included), N = 4 samples per condition, mean ± SEM, Two-way ANOVA. P-values indicate comparisons to the qNSPC condition. (D, E) Circular bar plots based on LD-lipids quantification, depicting logarithmic fold changes (log2FC) of individual lipid species, with their number of carbon atoms: number of double bonds information shown for each lipid, in qNSPCs compared to prolNSPCs (D) and in qNSPCs compared to NSPC-derived astrocytes (E), color-coded by lipid class. Significantly up-(bars directed outside of the circle) or downregulated (bars directed inside the circle) lipids are shown with 100% opacity; bars with 50% opacity represent lipids without significant abundance changes. The p-adjusted cut-off is 0.05 (multiple-comparison adjusted). Lipid quantification is based on four biological replicates. PC: phosphatidylcholines, PE: phosphatidylethanolamines. (F, G) Heatmaps of TGs grouped by number of double bonds (F) or by number of carbon atoms (G), showing up-(yellow) or downregulated (purple) Z-scores in LD lipidomes of qNSPCs, prolNSPCs and NSPC-derived astrocytes.

We next analyzed the distribution of TGs and CEs from the LD core (Fig. 4C). Intriguingly, while qNSPCs LDs contained 85 mol% (of total lipid content, all detected lipid species included) TGs and 15 mol% CEs, prolNSPCs and astrocytes had significantly less TGs and more CEs in their LDs, shifting the ratio to ca. 40 mol% / 60 mol% TGs to CEs. These findings correlate with our observation that LDs in prolNSPCs and astrocytes appear smaller than those in qNSPCs, as CE-rich LDs have been associated with smaller LD sizes (*35*, *36*). Despite differences in LD-core composition, the LD-monolayer phospholipid distribution in qNSPCs and astrocytes was very similar: 1.01 mol% PC and 0.25 mol% PE in qNSPCs, and 1.03 mol% PC and 0.17 mol% PE in astrocytes (fig. S4B). In contrast, prolNSPC LDs had much lower PC and PE levels: 0.57 mol% and 0.066 mol%, respectively.

To further understand LD-lipidome differences, we performed pairwise comparisons of mol% of detected lipid species between cell types, pooling TG species with the same total carbon number and double-bound number for simplicity (Fig. 4D, E, fig. S4C). The strongest differences appeared between qNSPCs and prolNSPCs: most TGs and phospholipids were significantly upregulated in qNSPCs, while two of four CE species were significantly downregulated (Fig. 4D). LD lipids of qNSPCs differed less from those of astrocytes, showing significant upregulation of some TGs with fewer carbon atoms and two PEs, and significant downregulation of CE 17:1 (Fig. 4E). Comparison of astrocytic and prolNSPC LD lipidomes showed significant upregulation of several phospholipids and some polyunsaturated TGs with higher carbon numbers (fig. S4C). Analysis of the relative amounts of the ten most abundant TG species and all detected CE, PC, and PE lipids further illustrates these shifts (fig. S4D-G).

Saturation levels and carbon numbers of TGs also differed between groups: prolNSPC LDs showed overall downregulation of all TG species and had fewer double bonds and carbon atoms; qNSPCs accumulated polyunsaturated TGs with intermediate carbon numbers; astrocytic LDs were enriched in highly polyunsaturated TGs with higher carbon numbers (Fig. 4F-G). Interestingly, qNSPC LDs were depleted in saturated TGs, consistent with studies showing that saturated TGs lead to smaller artificial LDs (*37*).

Concluding, we established an endogenous LD lipidome for NSPCs and their progeny, whose composition is highly cell-type dependent.

### CIDE proteins are involved in LD size regulation of qNSPCs

We have provided microscopic (Fig. 1B, C) and molecular evidence (Fig. 2-4) that LDs undergo strong structural remodeling upon changes in NSPC state. However, whether such LD changes play a role in NSPC function remains unclear. Our proteomic analysis showed that among the most upregulated LD proteins in qNSPCs were two CIDE-family members – CIDEA and CIDEB (Fig. 5A). Moreover, these proteins were exclusively found on qNSPC LDs, with no signal in LD fractions and total proteome fractions of prolNSPCs or astrocytes (table S4). The primary function of CIDE proteins is to control LD fusion and growth (Fig. 5B) (*11*). Thus, their exclusive presence on qNSPC LDs could theoretically explain the large LDs in this cell type.

**Fig. 5.**
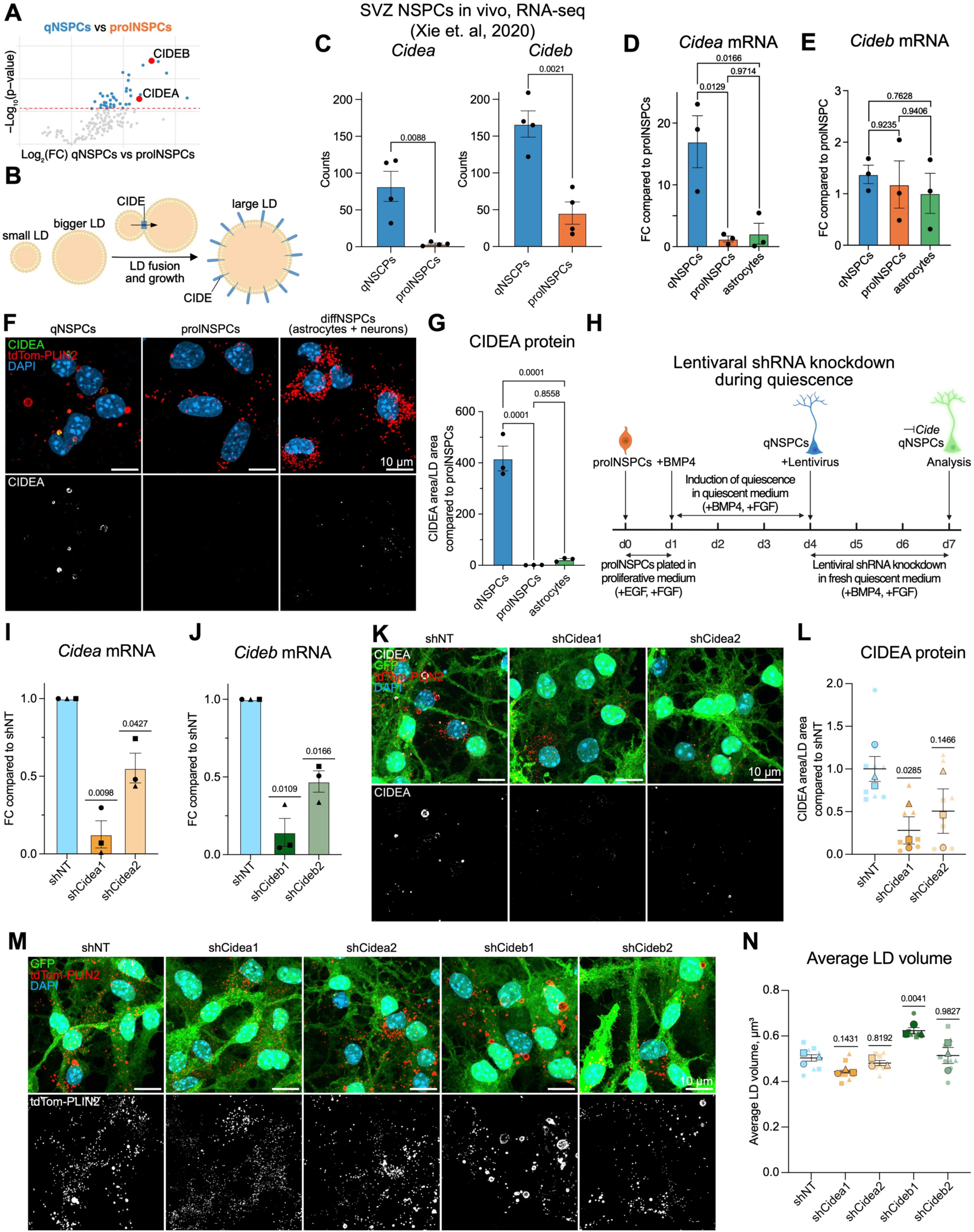
Knockdown of qNSPC-specific LD proteins CIDEA and CIDEB affects LD size. (A) Volcano plot based on high-confidence lipid droplet (LD) protein quantification in quiescent neural stem/progenitor cells (qNSPCs) compared to proliferative NSPCs (prolNSPCs), highlighting CIDEA and CIDEB as significantly upregulated on LDs in qNSPCs. (B) Illustration of CIDE-mediated LD growth by fusion, in which CIDE proteins serve as structural units of a molecular tunnel created between LDs to enable lipid flux. (C) *Cidea* (left) and *Cideb* (right) mRNA counts in qNSPCs compared to prolNSPCs based on bulk RNA-seq data from murine SVZ NSPCs in vivo collected by Xie and colleagues (*45*). (D, E) Analysis of *Cidea* (D) and *Cideb* (E) mRNA expression by qRT-PCR in wild-type qNSPCs, prolNSPCs and NSPC-derived astrocytes), normalized to *beta*-*actin* expression. Each dot represents an independent experiment, fold change (FC) relative to prolNSPC *Cide* levels ± SEM, one-way ANOVA. (F) Representative confocal images of qNSPCs, prolNSPCs, and differentiating NSPCs (diffNSPCs, co-culture of astrocytes and neurons) derived from the tdTom-Plin2+/- mice, showing endogenous CIDEA protein levels. Representative images are maximum-intensity projections. (CIDEA: CIDEA protein, tdTom-PLIN2: LDs, DAPI: nuclei). (G) Quantification of CIDEA protein levels in wild-type qNSPCs, prolNSPCs and NSPC-derived astrocytes. Each dot represents the average of images coming from the same coverslip, fold change (FC) relative to prolNSPC CIDE levels ± SEM, one-way ANOVA. (H) Schematic of the experimental design for lentiviral short-hairpin RNA (shRNA) knockdown (KD) of *Cide* mRNAs during quiescence. EGF: epidermal growth factor, FGF: fibroblast growth factor, BMP4: bone morphogenic protein 4. (I, J) Analysis of *Cidea* (I) and *Cideb* (J) mRNA expression by qRT-PCR in qNSPCs from tdTom-Plin2+/- mice upon *Cidea* (I) or *Cideb* (J) KD using two different shRNAs. mRNA levels are normalized to *beta*-*actin*. Each dot of the same shape represents an independent experiment, FC compared to control non-targeting shRNA (shNT) ± SEM, one sample t- and Wilcoxon-test. (K) Confocal images of qNSPCs from tdTom-Plin2+/- mice showing CIDEA protein levels upon *Cidea* KD using two different shRNAs. Representative images are maximum-intensity projections. (CIDEA: CIDEA protein, tdTom-PLIN2: LDs, GFP: GFP reporter co-expressed from shRNA plasmid, DAPI: nuclei). (L) Quantification of CIDEA protein levels upon *Cidea* KD using two shRNAs. Each large dot of the same shape represents the average of one independent experiment; each small dot represents the average of images coming from the same coverslip. N = 3, n = 9. Mean ± SEM of N = 3, one-way ANOVA. (M) Representative confocal images of qNSPCs from tdTom-Plin2+/- mice showing LD phenotype upon *Cidea* and *Cideb* KD using two shRNAs. Representative images are maximum-intensity projections. (tdTom-PLIN2: LDs, GFP: GFP reporter co-expressed from shRNA plasmid, DAPI: nuclei). (N) Quantification of average LD volumes upon *Cidea* and *Cideb* KD using two shRNAs. Each large dot of the same shape represents the average of one independent experiment; each small dot represents the average of images from the same coverslip. N = 3, n = 9. Mean ± SEM of N = 3, one-way ANOVA.

CIDE proteins show tissue- and cell type-specific distribution: CIDEA is abundant in brown adipocytes, and CIDEB in hepatocytes (*38*, *39*). In the mouse brain, *Cidea* and *Cideb* expression is near detection limits in cortex and hippocampal cells, including DG (*40–44*). However, by exploring the RNA-seq dataset from the study focusing on in vivo adult mouse SVZ stem-cell niche by Xie and colleagues (*45*), we found that *Cidea* and *Cideb* are specifically expressed at low levels in qNSPCs, but not in prolNSPCs, (Fig. 5C).

To further explore CIDE function in NSPC quiescence, we quantified by qRT-PCR *Cidea* and *Cideb* messenger-RNA (mRNA) levels in WT qNSPCs, prolNSPCs, and astrocytes (Fig. 5D, E). qNSPCs showed ∼17-fold upregulation of *Cidea* mRNA levels compared to prolNSPCs and astrocytes (Fig. 5D). Surprisingly, *Cideb* mRNA levels were equal across all three cell types (Fig. 5E), despite its exclusive protein presence on qNSPC LDs, indicating that the protein levels are likely not regulated only at the transcriptional level. To further assess protein levels of CIDE family members with immunocytochemistry we used primary tdTom-Plin2 NSPCs (*18*). While we did not find commercial CIDEB antibody that efficiently detected CIDEB protein in murine NSPCs, we successfully validated that CIDEA protein is present on qNSPC LDs but absent from prolNSPC and diffNSPC LDs (Fig. 5F-G).

To manipulate *Cide* levels in qNSPCs, we designed a lentiviral short-hairpin RNA (shRNA) knockdown (KD) experiment, in which tdTom-Plin2 qNSPCs were infected after three days in quiescent medium with lentivirus carrying shRNA plasmids with a GFP reporter to knock down *Cidea* or *Cideb* (Fig. 5H). Cells remained in virus-containing medium for three additional days (six total days of quiescence) before analysis. We used two shRNAs per gene (shCidea1/2, shCideb1/2) and a non-targeting control (shNT). qRT-PCR analysis showed an 87% reduction in *Cidea* mRNA with shCidea1 and 45% with shCidea2, while *Cideb* mRNA remained unchanged upon *Cidea* KD (Fig. 5I, fig. S5A). Conversely, *Cideb* KD reduced *Cideb* mRNA by 86% (shCideb1) and 53% (shCideb2) (Fig. 5J), with *Cidea* expression unaffected (fig. S5B). Immunocytochemistry showed a 72% reduction in CIDEA protein with shCidea1 and 50% with shCidea2, with no change upon *Cideb* KD (Fig. 5K, L, fig. S5C, D). Together, we confirmed efficient *Cidea*/CIDEA KD on mRNA and protein levels and *Cideb* KD on mRNA level.

Because CIDE proteins mediate LD growth via fusion, we hypothesized that *Cide* KD would reduce LD size in qNSPCs. Surprisingly, although *Cidea* KD tended to reduce LD size compared to shNT, the change was not significant; total LD volume/cell remained unchanged, and LD number/cell tended to increase (Fig. 5M, N, fig. S5E-G). More importantly, *Cideb* KD produced the opposite effect: average LD volume significantly increased, total LD volume/cell remained unchanged, and LD number/cell tended to drop, suggesting an additional role of CIDEB beyond LD fusion (Fig. 5M, N, fig. S5E-G). The second shRNAs (Cidea2, Cideb2) did not significantly alter LD phenotype, likely due to milder KD efficiency.

### CIDEB KD leads to transcriptional features resembling a senescence-like transition in qNSPCs

To further understand the functional role of CIDE proteins in qNSPCs, we performed bulk RNA barcoding and sequencing (BRB-seq) in qNSPCs with lentiviral *Cide* KD (tables S6). Both *Cidea* and *Cideb* mRNA levels were significantly downregulated, confirming KD efficiency; however, as observed before, the second shRNAs were less efficient for both genes (Fig. 6A, B, fig. S6A, B). KD of *Cidea* had very little effect on the transcriptome, with only 1-2 genes showing significant expression changes (fig. S6C, D, table S7), suggesting that altering *Cidea* does not affect the qNSPC transcriptome. In contrast, depletion of *Cideb* with shCideb1 caused massive transcriptomic changes, with many genes significantly up- and downregulated (Fig. 6C, table S7). In line with the milder KD efficiency and LD phenotype of shCideb2, these changes did not reach statistical significance under the shCideb2 condition when using the same filtering criteria (Fig. 6D, table S7). However, color-coding the shCideb2 volcano plot based on shCideb1 differentially expressed genes showed that significantly downregulated genes in shCideb1 also had negative log2FC in shCideb2 condition, and significantly upregulated genes had positive log2FC (Fig. 6E), indicating a similar but weaker transcriptomic remodeling.

**Fig. 6.**
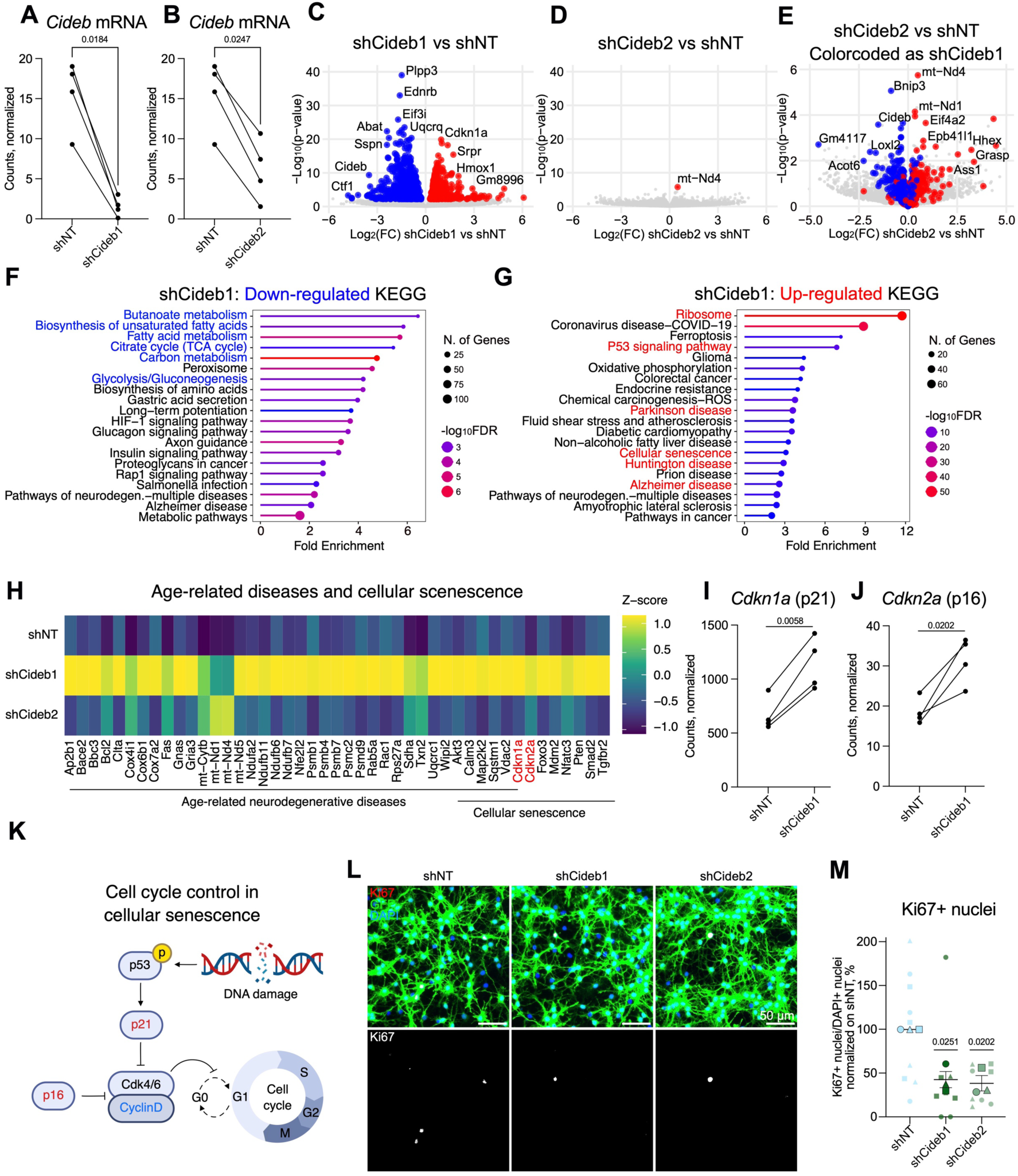
Bulk RNA-seq reveals transcriptomic remodeling of qNSPCs toward a senescence-like phenotype upon CIDEB depletion. (A, B) Levels of *Cideb* mRNA in quiescent neural stem/progenitor cells (qNSPCs) upon the *Cideb* knockdown (KD) with shCideb1 (A) or shCideb2 (B) compared to non-targeting shRNA (shNT), based on bulk RNA barcoding and sequencing (BRB-seq). Each dot represents an independent experiment, with dots from the same experiment connected by a line. The same shNT control is shown in panels A and B, as both KD experiments were performed in parallel and therefore share a common control. KD significance was tested using a ratio paired t-test. (C, D) Volcano plots based on BRB-seq quantification showing gene-expression changes in qNSPCs upon *Cideb* KD with shCideb1 (C) or shCideb2 (D) compared to shNT. Significantly upregulated genes are highlighted red, significantly downregulated genes in blue. Gray dots represent genes without significant changes. p-adjusted cut-off is 0.05 (multiple-comparison adjusted); fold change cut-off is 1.2. Quantification is based on four biological replicates. (E) Volcano plot showing gene-expression changes in qNSPCs upon *Cideb* KD with shCideb2 compared to shNT. Genes significantly upregulated in the shCideb1 condition are highlighted in red, significantly downregulated genes in blue. Gray dots represent genes not significantly changed in shCideb1. Quantification is based on four biological replicates. (F, G) Kyoto Encyclopedia of Genes and Genomes (KEGG) on ontology analysis of downregulated (F) and upregulated (G) genes upon *Cideb* KD with shCideb1 in qNSPCs. Selected downregulated pathways are highlighted blue, selected upregulated pathways are highlighted red. (H) Heatmap of genes belonging to age-related neurodegenerative diseases and cellular senescence pathways from (G), showing relative expression (Z-score) in qNSPCs upon *Cideb* KD with shCideb1, shCideb2, and shNT. Genes involved in senescence-associated cell-cycle regulation Cyclin Dependent Kinase Inhibitor 1A and 2A (*Cdkn1a* and *Cdkn2a*) are highlighted in red. (I, J) Levels of *Cdkn1a* (p21) (I) and *Cdkn2a* (p16) (J) mRNA in qNSPCs upon *Cideb* KD with shCideb1 compared to shNT, based on the BRB-seq. Each dot represents an independent experiment, with dots from the same experiment connected by a line. Significance tested using a ratio paired t-test. (K) Schematic representation of cell-cycle control during cellular senescence. Genes significantly upregulated in BRB-seq analysis are highlighted in red, significantly downregulated genes in blue. (L) Single-plane epifluorescence images of qNSPCs show Ki67 protein levels upon *Cideb* KD using two shRNAs. (Ki67: Ki67 protein, GFP: GFP reporter co-expressed from shRNA plasmid, DAPI: nuclei). (M) Quantification of Ki67+ nuclei upon *Cideb* KD using two shRNAs. Each large dot of the same shape represents the average of one independent experiment; each small dot represents the average of images from the same coverslip. N = 3, n = 9. Mean ± SEM of N = 3, one sample t- and Wilcoxon-test.

Gene-ontology analysis using Kyoto Encyclopedia of Genes and Genomes (KEGG) revealed fatty-acid and glucose metabolism among the top downregulated pathways in qNSPCs with *Cideb* KD (Fig. 6F, table S8). Among the top upregulated pathways were ribosomal biogenesis, p53 signaling, and pathways involved in cellular senescence and age-related neurodegenerative diseases such as Parkinsońs, Huntingtońs and Alzheimeŕs (Fig. 6G, table S8).

Given that genes involved in the p53 pathway, which plays a critical role in senescence and aging (*46*), as well as genes linked to neurodegenerative diseases were upregulated upon *Cideb* KD, we next examined the genes within these gene ontology terms. Compared to control shRNA, shCideb1 qNSPCs showed significant upregulation of *Cdkn1a* (p21), a key player in p53-induced senescence whose induction leads to cell-cycle arrest (Fig. 6H, I). Moreover, *Cdkn2a* (p16), which is a central regulator of senescence-induced cell-cycle arrest independent of p53, was also significantly upregulated (Fig. 6H, J). Furthermore, many genes in the age-related neurodegeneration category upregulated upon *Cideb* KD pointed toward mitochondrial/oxidative-stress responses (Fig. 6H). Although less pronounced, many of these genes were also mildly upregulated with shCideb2 (Fig. 6H). Consistent with these observations, Ki67 signal, which is a marker for actively cycling cells and low in quiescent NSPCs, was even further reduced in shCideb1/2 conditions, in line with a senescence like cell-cycle arrest (Fig. 6K-M). Importantly, we found that *Cideb* mRNA expression declines with aging in qNSPCs in vivo in a study examining age-related transcriptomic changes in mouse NSPCs, further supporting a link between *Cideb* levels and a senescence-like phenotype (fig. S6E) (*45*).

Together, these results show extensive transcriptomic remodeling upon *Cideb* depletion and suggest a potential role for CIDEB in maintaining a healthy qNSPC state, with its loss inducing a senescence-like phenotype.

## Discussion

The maintenance of NSPC quiescence, which preserves the long-term capacity of the NSPC pool, and the regulation of activation processes that allow NSPC expansion for subsequent differentiation, is essential for maintaining adult neurogenesis. Numerous intrinsic and extrinsic factors regulate NSPC behavior, among which cellular metabolism has recently been recognized as a key determinant of NSPC fate. Notably, in hippocampal neurogenesis, lipid metabolism controls prolNSPC proliferation, with FAO maintaining NSPC stemness, whereas lipogenesis drives differentiation (*8*, *47*, *48*). However, we lack a mechanistic understanding of how lipid metabolism regulates NSPC behavior.

In this study, we generated the first comprehensive endogenous LD proteome and lipidome of NSPCs and their progeny, allowing us to reveal striking differences in LD composition across NSPC states. Using this dataset, we identified CIDEB, a protein specifically enriched on qNSPC LDs, whose KD was associated with altered LD size and transcriptional remodeling towards a senescence-like phenotype, suggesting a role in quiescence maintenance.

Establishing a high-confidence LD proteome is challenging. LDs are highly interactive organelles and form MCSs with most other organelles, which can lead to co-purification of non-LD proteins even in highly pure LD fractions. To overcome this, different groups have developed strategies to identify *bona fide* LD proteomes. Bersuker and colleagues employed APEX2-based proximity labelling to selectively tag LD proteins, enabling high-confidence proteome mapping in two human cell lines (*24*). Protein-correlation profiling across multiple organelles has identified high-confidence LD proteomes in *Drosophila* S2 cells and in murine liver under different metabolic conditions (*25*, *49*). Another study identified proteins strongly enriched on LDs in human macrophages and breast-cancer cells by comparing total-cell proteomes to LD-enriched fraction (*26*). While the proximity labelling physically tags LD proteins and thus provides high confidence in LD localization, it requires overexpression of the APEX system. As we aimed to study endogenous WT LDs, we choose the strategy of comparing LD fractions to total-cell proteomes to establish a high-confidence LD proteome of NSPCs and their progeny.

A further challenge in LD-proteome studies is the dependence of protein recruitment to the LD monolayer on LD-core lipid content and LD size (*50*). In fact, since many cell types do not form sufficient amount of LDs for extraction, LD formation is commonly induced by loading cells with oleic acid or acetylated apolipoprotein-B-containing lipoproteins (*24*, *26*). It is unknown whether the lipid content of artificially induced LDs corresponds to the lipids that LDs would naturally accumulate in the respective cell type, or whether this affects LD-coating protein composition.

Another level of complexity arises from the increasingly recognized LD heterogeneity between the cell types and even between LDs in the same cell type (*33*). While LDs share a common set of ubiquitously detected proteins (such as PLIN family proteins), many proteins localize to LD surface in a cell-types specific manner (*24*, *26*). Consequently, LD-proteome datasets from distinct cell types are needed to better understand cell-type-specific LD functions across tissues, including the brain. Thus, our endogenous LD proteome of NSPCs and their progeny contributes substantially to LD biology field and provides important insights into LD-protein specificity in NSPCs.

In this study, we analyzed the LD proteome of qNSPCs, prolNSPCs, and NSPC-derived astrocytes (Fig. 1–3, fig. S1–S3). Among the 171 high-confidence LD-associated proteins identified, NSPCs and their progeny shared a core of 42 LD proteins, 31 of which had previously been annotated as LD-associated (Fig. 2). Importantly, we also identified proteins that localized to LDs in a cell-type-specific manner, with the most pronounced differences in qNSPCs. This cell type exhibited a large number of LD proteins uniquely detected in qNSPCs across all functional categories. These observations align with current views in LD biology: LD proteomes share a common set of proteins but also display cell-type-specific heterogeneity (*24*, *26*). Notably, some classical LD-associated proteins consistently identified across datasets, such as RAB-family members, were detected exclusively on qNSPC LDs and not in the other two cell types (Fig. 3). This may reflect molecular-crowding effect, suggesting that RAB proteins preferentially associate with large and/or TG-rich LDs, underscoring the importance of studying endogenous LDs. At the same time, our dataset confirms recent studies mapping non-canonical LD proteins, such as APOE and TMEM263, to LD surface (*51*, *52*).

While LDs are central organelles in lipid metabolism, their lipidome remains far less characterized than their proteome. Although lipidomics is a rapidly developing field, significant technical challenges persist, from sample preparation to lipid species identification and data analysis (*53*). Much of our current knowledge of the LD lipidome derives from naturally lipid-rich tissues and cell types such as adipocytes, hepatocytes, or steatotic cells, and only a limited number of lipidomic datasets exist for isolated LDs (*35*). It is well established that the LD core is primarily composed of TGs and CEs, although additional lipid species have been reported at lower abundance (namely monoalk(en)yl diacylglycerols and fatty acid esters of hydroxy fatty acids in the LD core, and cholesterol, diacylglycerides, and ceramides at the neutral-core/monolayer interface) (*35*). In this study, we employed a targeted lipidomic approach covering a broad spectrum of lipid species. While this strategy limited identification of some lipid classes (such as diacylglycerols), it enabled quantification of the most classical LD core and monolayer species.

Our analysis revealed that LDs in qNSPCs exhibit a TG:CE ratio of approximately 85:15, which shifted markedly to 40:60 in prolNSPCs and NSPC-derived astrocytes (Fig. 4). This observation is consistent with growing evidence that LD size and protein composition are strongly influenced by LD-core lipid composition (*54–56*). Differences in TG and CE content may therefore underlie functional LD heterogeneity, potentially reflecting distinct LD-associated functions depending on LD size and lipid composition.

These findings underscore the importance of analyzing LDs with distinct lipid compositions in different cellular contexts and, if possible, without lipid loading to trigger LD formation. Notably, in our system, the medium was identical across all cell types (except for growth-factor addition or withdrawal), suggesting that the observed LD-lipidome differences reflect intrinsic lipid-handling differences rather than exogenous lipid sources. To date, lipidomic analyses of LDs in neural cells remain extremely limited, and only a few recent studies have begun to explore LD lipid composition in astrocytes or neurons upon exogenous oleic-acid loading or blockade of triglyceride lipolysis (*57*, *58*). These initial studies mark an underexplored research area and emphasize the substantial knowledge gap regarding LD lipid composition in neural cells.

Understanding LD structure across NSPC states is essential for gaining insight into NSPC biology and, in particular, the contribution of LDs to regulating NSPC states. In our study, we found many LD proteins specifically enriched in qNSPCs, suggesting they may be involved in maintaining or regulating quiescence. Interestingly, many proteins mediating organelle contact were specifically present on qNSPC LDs, suggesting that differential LD-organelle interactions might influence NSPC state. However, because LDs can interact with many organelles and quantitative analysis of contact sites requires high-resolution 3D imaging, further studies are needed to determine whether these LD-protein differences indeed reflect differential organelle contacts.

As a proof of concept that the cell-type-specific LD proteome is functionally relevant for NSPC behavior, we here focused on CIDE proteins, as they emerged as highly specific for qNSPCs compared to prolNSPCs and NSPC-derived astrocytes. Although CIDE proteins have not classically been associated with brain tissue, low RNA expression levels of *Cidea* and *Cideb* can nevertheless be detected in transcriptomic datasets of neurogenic niches (*45*, *59*). Importantly, these genes are selectively enriched in qNSPCs, but not in prolNSPCs or progenitors, in vivo, aligning with our findings (*45*, *59*). Their restricted and low-level expression in qNSPCs may explain why they have not been previously described as brain-expressed genes.

CIDE proteins are typically found in lipogenic tissues (adipose tissue, liver, intestines), where they regulate LD fusion, lipid secretion, and lipid synthesis (*11*). Thus, their specific expression in qNSPCs could underlie the LD phenotype we observe, characterized by a subset of large LDs. Interestingly, however, only depletion of *Cidea*, rather than *Cideb*, supported the expected LD-fusion phenotype, reflected by a slight but not significant LD size reduction. *Cideb* depletion instead caused abnormal LD enlargement, raising the possibility that CIDEB fulfills additional cellular functions beyond LD fusion. Together, our data indicate that CIDEA and CIDEB are specifically expressed in qNSPCs, but their precise roles remain unclear.

To investigate CIDE function in qNSPCs, we performed bulk RNA-seq. We found that CIDEB depletion triggers extensive transcriptional remodeling toward a senescence-like state, including upregulation of the p53 signaling pathway, increased expression of *Cdkn1a* (p21) and *Cdkn2a* (p16), two key regulators of G0 cell-cycle arrest, and reduced Ki67 protein expression, indicating impaired cell-cycle progression. Remarkably, analysis of published in vivo RNA-seq data showed that *Cideb* mRNA levels decline sharply in qNSPCs of aged mice compared to young animals (fig. S6E) (*45*). The same study reported upregulation of translational machinery and ribosomal subunits in aged qNSPCs, which were likewise strongly upregulated upon *Cideb* KD in our dataset. Furthermore, a recent CRISPR-Cas9 screen by Ruetz and colleagues identified regulators of NSPC aging (*60*). Notably, *Cideb* knockout inhibited activation of old qNSPCs 14 days after induction of proliferation in vitro (fig. S6F). These findings support our observation that CIDEB depletion induces a senescence-like phenotype and suggest a potential role for CIDEB in maintaining a healthy quiescent state.

While these data point toward an important function of CIDEB in qNSPCs, additional functional experiments are required to directly link *Cideb* expression levels to qNSPC fitness. Further studies must clarify how *Cideb* depletion drives senescence-like phenotype and how this relates to LD biology. Notably, our study examined LD structure in NSPCs isolated from the SVZ of male mice. Whether LD architecture is conserved across NSPC states in the DG, another neurogenic niche in the mouse brain, remains unknown. Moreover, it will be important to determine whether LD content in NSPCs is sex-dependent by analyzing LD structure in female-derived NSPCs.

In conclusion, our study provides the first comprehensive endogenous LD proteome and lipidome of NSPCs and their progeny, revealing strong cell-state specificity in LD composition and establishing a valuable resource for investigating LD biology in NSPC maintenance and lineage progression.

## Materials and Methods

### Animals

Male C57BL/6J mice were obtained from Janvier Labs (France). Heterozygous tdTom-Plin2 mice on a C57BL/6J background were maintained through in-house breeding as previously described (*18*). Animals were housed in individually ventilated cages with ad libitum food and water. Housing conditions were dark/light cycle 12/12, ambient temperature around 21-22 °C and humidity between 40 and 70% (55% in average). All experiments including animals were approved in accordance with the Swiss law after approval from the local authorities (Cantonal veterinary office, Canton de Vaud, Switzerland). NSPCs were isolated from SVZs of 8-week-old male mice of the respective strains as described below.

### Cell culture

Adult mouse NSPCs were isolated from the SVZ of 8-week-old C57BL/6J and tdTom-Plin2+/- male mice as described previously (*9*). In short, mice were briefly anaesthetized with isoflurane before decapitation. The SVZs were micro-dissected, and the tissue was dissociated into a single-cell suspension using the GentleMacs Dissociator (Milteny) and the papain-based MACS Neural Tissue Dissociation Kit (Milteny, #130-092-628), following the manufacturer’s instructions. Myelin debris was then removed using the MACS myelin removal beads (Milteny, #130-096-731) and a QuadroMACS Separator (Milteny, #130-090-976). Isolated single-cells were cultured as neurospheres in DMEM/F12/GlutaMAX (Invitrogen, #31331-028) supplemented with B27 (Invitrogen, #17504044), 20 ng/ml human EGF (AF-100-15, PeproTech), 20 ng/ml human basic FGF-2 (100-18B, PeproTech), and 1× PSF (Invitrogen, #15240062) at 37 °C and 5% CO_2_. Medium was changed every 2-3 days. The neurospheres were passaged at least 5 times before experimental use to remove progenitors or other proliferating cells. Subsequently, cells were adapted to N2-supplemented medium instead of B27 (proliferation condition). NSPCs isolated from 3-4 mice were pooled for each preparation, and experiments were performed using cultures up to passage 25. To propagate NSPCs, cells were maintained in uncoated standard plastic cell culture dishes (Corning, #430167, 100 × 20 mm, TC-treated). For all experimental assays, glass coverslips (10337423, Fisher) or standard plastic cell culture dishes were coated with 50 µg/ml (glass) or 10 µg/ml (plastic) poly-L-ornithine (Sigma, Cat. #P3655) and 5 µg/ml laminin (Sigma, #L2020-1MG) before cell seeding.

#### Proliferation condition

ProlNSPCs were maintained in DMEM/F12/GlutaMAX (Invitrogen, #31331-028) complemented with N2 (17502048, Gibco), 20 ng/ml human EGF (AF-100-15, PeproTech), 20 ng/ml human basic FGF-2 (100-18B, PeproTech), 5 µg/ml Heparin (H3149-50KU, Sigma) and 1× PSF (Invitrogen, #15240062) (*9*). Culture medium was replaced every 2-3 days. For experiments, cells were seeded at a density of 30 000 cells/cm^2^ onto coated culture dishes of glass coverslips.

#### Quiescence induction

Quiescence was induced as previously described (*61*). Briefly, NSPCs were seeded at 47 000 cells/cm^2^ on coated plates or glass coverslips and medium was replaced with quiescence medium containing DMEM/F12/GlutaMAX, N2, 20 ng/ml human basic FGF-2, 5 µg/ml Heparin, 50 ng/ml BMP4 (RnD Systems #5020-BP) and 1× PSF after 24 h. Cells reached a fully quiescent state 72 h after quiescence induction.

#### Differentiation condition

prolNSPCs were plated at 100 000 cells/cm^2^ on coated flasks or glass coverslips. Cells were initially cultured in differentiation medium containing reduced amounts of growth factors. (The same medium as for proliferation condition, but only 1/5th of EGF and bFGF2 in first two days and no EGF and bFGF2 after). The medium was fully changed after 2 days to medium without EGF and bFGF2. Cells were fixed or collected after 5 days of differentiation.

#### Lentiviral shRNA KD during quiescence

Quiescence in NSPCs was induced as stated above, until the cells reached fully quiescent state after 72 h in quiescent medium. The medium was then exchanged to a fresh quiescent medium that contained 1 µl of high-titer lentiviral solution per 1 ml of cell culture medium (infection ratio achieved with this amount of virus was 92.3% ± 5.2%). qNSPCs were incubated with the virus at 37 °C, 5% CO2 for another 72 h before being fixed or collected.

#### HEK293T cell culture

HEK293T cells were cultured in DMEM/high glucose/GlutaMAX (Invitrogen, # 61965-026) complemented with 10% fetal bovine serum (Thermo Fisher Scientific, A3840101) at 37 °C (5% CO2). The cells were regularly split to keep them sub-confluent. To achieve this, culture medium was removed from the standard plastic cell culture dishes (Corning, #430167, 100 × 20 mm, TC-treated), the cells were washed with 10 ml of PBS (1X) and dissociated from the flask surface by the addition of 2 ml TripLE^TM^ Express enzyme 1X (Fisher Scientific, 12604-013-100ML). The reaction was stopped when 2 ml of culture medium were added to the dissociated cells. After that, the cell suspension was diluted and transferred to the new cell culture dish.

### Immunocytochemistry

Cells were fixed with 37 °C paraformaldehyde/PBS (4%) for 20 min at room temperature (RT), washed 3 x with 1x PBS for 10 min and were subsequently stored at 4 °C.

#### TdTom-PLIN2-based LD visualization

For the LD morphology analysis in tdTom-PLIN2+/- NSPCs, the endogenous tdTomato signal was used without cell permeabilization. Following fixation, nuclei were counterstained with DAPI (D9542, Sigma) diluted in 1× TBS for 10 min. Coverslips were subsequently washed twice with 1× TBS for 5 min and mounted using a homemade PVA-DABCO-based mounting medium.

#### Saponin-based staining protocol

Immunostaining procedures requiring simultaneous visualization of LDs visualization were performed using a saponin-based protocol as previously described (*62*). Briefly, cells were blocked with blocking buffer (1.5% Glycine, 3% BSA, 0.01% Saponin in 1× PBS) for 45 min, RT. Primary antibodies were diluted in antibody diluent (0.1% BSA, 0.01% Saponin in 1× PBS) and incubated with the samples overnight at 4 °C. Anti-CIDEA antibody (1:250, Proteintech, 13170-1-AP) was used with saponin-based protocol. After three 10 min washes with 1× PBS, cells were incubated for at least 1 h at RT with appropriate secondary antibody diluted in antibody diluent, while protected from light. Following a 10 min wash with 1× PBS, nuclei were stained with DAPI (D9542, Sigma) diluted in 1× TBS for 10 min, followed by one wash in 1× TBS for 10 min. Coverslips were mounted in homemade PVA-DABCO-based mounting medium.

#### Triton-based staining protocol

For immunostaining experiments not involving LD visualization, cells were permeabilized and blocked in blocking buffer (0.25% Triton X-100, 3% donkey serum in 1× TBS) for 30 min at RT followed by incubation with primary antibodies. Primary antibodies were prepared in the same blocking solution and incubated with the cells overnight at 4 °C. Following primary antibodies were used with triton-based protocol: anti-Ki67 (1:500, Abcam, ab15580) and anti-GFP (1:2 000, Abcam, ab13970). Following three washes in 1x TBS (10 min each), cells were incubated with the corresponding secondary antibodies for at least 1 h at RT in the dark. Samples were then washed twice with TBS, counterstained with DAPI in 1× TBS for 10 min, and washed once more before mounting in homemade PVA-DABCO mounting medium.

### Image acquisition and analyses

All confocal images were acquired with a confocal microscope (Zeiss, LSM 900) with a 63x objective. Imaging of isolated LD fractions and imaging of Ki67+ nuclei was performed on an epifluorescence microscope (Nikon 90i) using a 20x and 40x objective. Acquisition parameters were held constant across states and replicates within each experiment.

#### LD morphology analysis

3-D LD reconstruction and characterization were done with the imaging software Imaris (https://imaris.oxinst.com/) using the “Surface” modules of ImarisCell. LD surfaces were reconstructed based on tdTom-PLIN2 signal using machine learning segmentation, whereby the machine learning system was trained on different images within an experiment and then the same training was applied to all images. The number of cells was quantified through manual counting of DAPI-positive nuclei.

#### CIDEA expression quantification

Quantification of CIDEA signal in NSPCs was done with Fiji (*63*). In brief, CIDEA acquisition was preprocessed using “Z-project” and tdTom-PLIN2 acquisition using “Z-project” and “Subtract Background”. CIDEA signal was converted to a mask after setting a fixed threshold (equal across all images) and tdTom-PLIN2 signal threshold was determined automatically with the “Moment” algorithm. Using the “Image Calculator” tool, thresholded CIDEA signal was masked on thresholded tdTom-PLIN2 signal to calculate CIDEA area that overlapped with tdTomPLIN2-positive LDs. Area covered by masked on tdTomPLIN2 CIDEA signal and area covered by tdTomPLIN2 signal only were quantified using “Analyze particles”. CIDEA area covered was then normalized to the total area of tdTomPLIN2.

#### Ki67 quantification

Quantification of Ki67 nuclei was done with Fiji. In brief, Ki67 and DAPI acquisitions were preprocessed using “Gaussian Blur” and “Set threshold” and converted to mask. The masks were processed with “Watershed” and number of nuclei in each channel was quantified using “Analyze Particles”.

### Cell collection for LD isolation

For LD isolation, NSPC cell culture was upscaled to ca. 100 million cells and collected as following:

#### prolNSPCs

Per LD isolation, cells were cultured on four standard plastic cell culture dishes (Corning, 430599, 150 × 25 mm, TC-treated) coated with poly-L-ornithine (Sigma, Cat. #P3655) and laminin (Sigma, #L2020-1MG) as described above until confluent. Cells were then washed down with 1 000 µl pipette in the old medium and collected in a pellet by centrifugation at 300 g for 3 min. The cell pellet was then resuspended in 50 ml of 1 x PBS following by the centrifugation at 300 g for 3 min and washed pellet was used for the subsequent LD isolation. Four independent LD isolations were performed.

#### qNSPCs

Per LD isolation, qNSPCs were cultured on four standard plastic cell culture dishes (Corning, 430599, 150 × 25 mm, TC-treated) coated with poly-L-ornithine (Sigma, Cat. #P3655) and laminin (Sigma, #L2020-1MG) as described above until reaching full quiescence (72 h in quiescent medium). For the cell collection, the old medium was aspirated from the plates, and the cells were washed with 1x PBS. To dissociate cells from the plate, they were treated with TripLE^TM^ Express enzyme 1X (Fisher Scientific, 12604-013-100ML) for up to 5 min. The enzyme was then inhibited by dilution with 1x PBS and cells were collected in the tube for the centrifugation at 400 g for 10 min at RT. The resulting pellet was used for the subsequent LD isolation. Four independent LD isolations were performed.

#### NSPC-derived astrocytes

For one LD isolation, diffNSPCs were cultured on ten standard plastic cell culture flasks (Corning, T175, 430823) coated with poly-L-ornithine (Sigma, Cat. #P3655) and laminin (Sigma, #L2020-1MG) as described above until reaching day 5 of differentiation. To isolate astrocytes from the neurons, flasks with the diffNSPCs were shaken for 1 h at 200 rpm, 37 °C and 5% CO_2_, whereby neurons detached from the flask bottom and astrocytes remained attached. The old medium was then aspirated from the plates, and the astrocytes were washed with 1x PBS. To dissociate cells from the plate, they were treated with TripLE^TM^ Express enzyme 1X (Fisher Scientific, 12604-013-100ML) for up to 10 min at 37 °C. The enzyme was then inhibited by dilution with 1x PBS and cells were collected in the tube for the centrifugation at 400 g for 10 min. The resulting pellet was used for the subsequent LD isolation. Four independent LD isolations were performed.

### LD isolation by ultracentrifugation

The cell pellet collected for the LD isolation was resuspended in 2.5 ml of ice-cold hypotonic lysis medium (HLM, 20 mM Tris-Cl pH 7.4, 1 mM EDTA pH 8.0) containing cOmplete, Mini, EDTA-free Protease Inhibitor Cocktail (Sigma-Aldrich, 11873580001) and incubated on ice for 10 min. Cells were lysed in nitrogen cavitation (Parr, 4639, 45 ml) at 35 bars for 15 min while on ice and collected lysate was centrifuged at 1000 g, 4 °C for 10 min. The supernatant was subsequently transferred to an Ultra-Clear centrifuge tube (Beckman-Coulter, 344060), resuspended in sucrose/HLM to a final concentration of 20% sucrose/HLM, and overlaid by 5 ml of 5% sucrose/HLM followed by 5 ml of HLM. The sample was then centrifuged at 28 000 g, 4 °C for 30 min in an ultracentrifuge using an SW41 swinging bucket rotor (Beckman-Coulter, 331362). LD fraction I (LDI) was collected using a tube cutter (Beckman-Coulter, 19175), and transferred to a clean Ultra-Clear centrifuge tube (Beckman-Coulter, 344060) for repeated centrifugation in the identical sucrose gradient at 28 000 g, 4 °C for 15 min. LD fraction II (LDII) was then collected in the same manner as LDI and used for LC-MS analysis.

### Western Blot

Proteins from the cytosolic fraction (CF) and LD fractions were precipitated in 8.17 volumes of 100% acetone (Sigma-Aldrich, 270725-1L) precooled to -20 °C and 1.5 volumes of 100% ice-cold trichloroacetic acid (Sigma-Aldrich, 91228-100G) at -20 °C overnight. The samples were then centrifuged at 17 200 g, 4 °C for 15 min and the protein pellet was washed three times in 1 ml of 100% acetone precooled to -20 °C by resuspending it by vortexing and repeating centrifugation step at 17 200 g, 4 °C for 15 min. After the final wash, the pellet was air-dried for 1 h. The protein pellet was subsequently resuspended in 30 µl of freshly prepared 8 M Urea with 4% SDS and incubated in this solution for 1 h at RT, vortexing every 5 min. Protein concentration was then measured with the Qubit Protein Assay Kit (Thermo Fisher Scientific, Q33211). Protein extracts were mixed with Laemmli sample buffer (#1610747, Bio-rad) supplemented with beta-mercaptoethanol and separated by SDS-PAGE on 12% PAA precast gel (#4561044, Bio-Rad). The gel was run at 50 V for 40 min and then at 100 V. The proteins were then transferred to an AmershamTM Protran® Western-Blotting-Membrane (#GE10600044, Sigma-Aldrich) at 110 V for 90 min. The membrane was then blocked in a blocking solution (5% milk powder, 0.05% Triton in 1x TBS) before incubation with primary antibody diluted in the blocking solution overnight at 4 °C. Following three washes (10 min each) in 1x TBS, membranes were incubated with secondary antibody in blocking buffer. Following three further washes (10 min each) in 1x TBS, proteins were revealed using ECL technology with WesternBright ECL HRP Substrate (K-12045-D20, Witec). The following antibodies were used: anti-Perilipin 2 rabbit antibody (1:1 000, Abcam ab52356), anti-β-Actin mouse antibody (1:5 000, Sigma-Aldrich A2228), peroxidase AffiniPure donkey anti-rabbit IgG (1:5 000, Jackson ImmunoResearch 711035-152), peroxidase AffiniPure donkey anti-mouse IgG (1:5 000, Jackson ImmunoResearch, 715-035-150).

### Proteomic analysis of LD fractions

#### Sample preparation and protein digestion

Samples were digested following a modified version of the in-StageTip (iST) method named miST method (*64*). For total proteome samples, cell pellets were lysed in miST lysis buffer (1% Sodium deoxycholate, 30 mM Tris pH 8.6, 10 mM DTT) and heated for 10 min at 75 °C. Samples were then diluted 1:1 (v:v) with water containing 4 mM magnesium chloride and benzonase (Merck, 70746, 100x dilution of stock = 250 Units/µl), and incubated for 15 min at RT to digest nucleic acids. Based on tryptophan fluorescence quantification, 25 µg of proteins were transferred to new tubes for digestion (*65*). Reduced disulfides were alkylated by adding ¼ vol. of 160 mM chloroacetamide (32 mM final concentration) and incubated for 45 min at RT in the dark. Samples were adjusted to 3 mM ethylenediaminetetraacetic acid (EDTA) and digested with 1.0 µg Trypsin/LysC mix (Promega, V5073) for 1 h at 37 °C, followed by a second 1 h digestion with an additional 0.5 µg of proteases.

For LD samples, after volume equilibration with 20 mM Tris pH 7.5, 100 µl of miST lysis buffer 2x were added to samples before heating them for 10 min at 75 °C. Reduced disulfides were alkylated by adding 16 mM (final concentration) of chloroacetamide and incubated for 45 min at RT in the dark. Samples were adjusted to 3 mM EDTA and digested with 2.0 µg Trypsin/LysC mix for 1 h at 37 °C, followed by a second 2 h digestion with an additional 1.0 µg of proteases.

To remove sodium deoxycholate, two sample volumes of isopropanol containing 1% trifluoroacetic acid (TFA) were added to the digests, and the samples were desalted on a strong cation exchange (SCX) plate (Oasis MCX 96-well, 186001830BA; WatersTM) by centrifugation. After washing with isopropanol/1% TFA and 2% acetonitrile/0.1% formic acid, peptides were eluted in 200 µL of 80% methyl cyanide, 19% H_2_O, and 1% (v/v) ammonia, then dried by centrifugal evaporation.

#### LC-MS/MS analyses

Data-dependent LC-MS/MS analyses of samples were carried out on a Fusion Tribrid Orbitrap mass spectrometer (Thermo Fisher Scientific) interfaced through a nano-electrospray ion source to an Ultimate 3000 RSLCnano HPLC system (Dionex). Peptides were separated on a reversed-phase custom packed 45 cm C18 column (75 μm ID, 100 Å, Reprosil Pur 1.9 µm particles, Dr. Maisch, Germany) with a 4%-90% acetonitrile gradient in 0.1% formic acid at a flow rate of 250 nl/min (total time 140 min). Full MS survey scans were performed at 120 000 resolution. A data-dependent acquisition method controlled by Xcalibur software (Thermo Fisher Scientific) was used that optimized the number of precursors selected (“top speed”) of charge 2+ to 5+ while maintaining a fixed scan cycle of 0.6 s. Peptides were fragmented by higher energy collision dissociation (HCD) with a normalized energy of 32%. The precursor isolation window used was 1.6 Th, and the MS2 scans were done in the ion trap. The m/z of fragmented precursors was then dynamically excluded from selection during 60 s.

#### Data processing

Data files were processed using MaxQuant (version 2.5.0.0) integrated with the Andromeda search engine (*66*, *67*). Carbamidomethylation of cysteine was set as a fixed modification, while methionine oxidation and protein N-terminal acetylation were specified as variable modifications. Database searches were performed against the mouse (*Mus musculus*) reference proteome from UniProt (www.uniprot.org; March 2023 release, comprising 55 309 sequences) and a contaminant database including common environmental contaminants and digestion-related enzymes (*68*). Mass tolerances were set to 4.5 ppm for precursor ions (after recalibration) and 0.5 Da for MS/MS fragments. Peptide and protein identifications were filtered at a 1% false discovery rate (FDR) based on matches to a decoy database generated by reversing protein sequences.

All subsequent analyses were done with an in house developed software tool (TARAM, available on https://github.com/UNIL-PAF/taram-backend). Contaminant proteins were removed, and IBAQ quantity values for protein groups were log2-transformed and samples were normalized using median subtraction methods (*69*). After assignment to groups, only proteins quantified in at least 3/4 samples of one group were kept. Missing values were imputed based on a normal distribution with a width of 0.3 standard deviations (SD), downshifted by 1.8 SD relative to the median. Student’s T-tests were carried out among conditions, with Benjamini-Hochberg correction for multiple testing (adjusted p-value threshold < 0.05). Imputed values were then removed before proceeding with further data analysis.

### Comparison of LD proteome of NSPCs and their progeny to the LD knowledge portal

To identify which proteins from our generated LD proteome of NSPCs and their progeny have already been associated with LDs, we have created a reference list of LD proteins by manually pooling the data from the studies used on the LD knowledge portal (*24–26*). Proteins that have been reported in at least one of these studies in at least one cell type or condition, were put in the final reference list, resulting in 310 proteins (table S3). The proteins from the LD reference list were then overlapped with our LD protein list of NSPCs and their progeny in R using the “%in%” operator.

### Lipidomic analysis of LD fractions

#### Sample Preparation

Each sample aliquot containing isolated LDs (50 µl) was extracted by adding isopropanol (250 µl), followed by vortex mixing for 30 s to ensure thorough homogenization. The extract was then transferred to 2.0 ml lysis tubes containing ceramic beads and homogenized using a Precellys homogenizer at 10 000 rpm for two cycles of 20 s each. The homogenates were subsequently centrifuged at 21 000 g for 15 min at 4 °C. An aliquot of the resulting supernatant (100 µl) was transferred to tubes containing pre-dried internal standards. The mixtures were vortexed for 30 s, sonicated for 3 min, and centrifuged again at 21 000 g for 15 min at 4 °C. The final supernatants were transferred to LC-MS vials equipped with glass inserts and analyzed using a TSQ Altis mass spectrometer (Thermo Fisher Scientific, USA), as described below.

#### Omics-scale targeted analysis of complex lipids (estimated quantification)

LD extracts were analyzed by hydrophilic interaction liquid chromatography coupled to electrospray ionization tandem mass spectrometry (HILIC-ESI-MS/MS) in both positive and negative ionization modes, using a TSQ Altis LC-MS/MS system (Thermo Scientific), as previously described by Medina et al. (*70*, *71*). The initial qualitative screen included 2 100 species (with transitions distributed across eight acquisition methods for optimal sensitivity and peak definition) belonging to five major lipid classes: glycolipids, cholesterol esters, sphingolipids, glycerophospholipids, and free fatty acids. Robustly detected species in pooled samples (representative of the entire batch but group-specific) were quantified across all samples, combining two runs – one in ESI positive mode and one in negative ionization mode. Chromatographic separation was carried out on an Acquity Premier BEH Amide column (1.7 µm, 100 mm × 2.1 mm I.D.; Waters, Milford, MA, USA) in a dual-column setup. The mobile phases consisted of: A = 10 mM ammonium acetate in acetonitrile:H₂O (95:5) (pH 8.2) and B = 10 mM ammonium acetate in acetonitrile:H₂O (50:50) (pH 7.4). The linear gradient elution was as follows: 0 min, 0.1% B; 2 min, 20% B; 5 min, 80% B; 8 min, 0.1% B; 12 min, 0.1% B. Following a 6-minute separation gradient in positive ionization mode, the first column was switched offline for conditioning while the second column was switched inline for a 6-minute separation in negative ionization mode, resulting in a total analysis time of 12 min per sample. The flow rate was 600 µL/min, the column temperature was 45 °C, and the injection volume was 2 µL. Optimized HESI source parameters were set as follows: voltage +3 500 V in positive mode and −2 500 V in negative mode; Sheath Gas = 60 (Arb), Aux Gas = 15 (Arb), Sweep Gas = 1 (Arb); Ion Transfer Tube Temperature = 380 °C. Nitrogen was used as the nebulizing gas and argon as the collision gas (1.5 mTorr). The vaporizer temperature was set to 350 °C.

#### Data Processing

Raw LC-MS/MS data were processed using the TraceFinder analysis software (Thermo Scientific). Estimated concentrations were calculated using single-point calibration with internal standards (75 IS at known concentrations from the Ultimate Splash mixture, Avanti Lipids). Quality assessment, including correction for signal-intensity drift, was performed using pooled quality-control (QC) samples analyzed periodically throughout the batch. Peaks showing high analytical variability, defined as a coefficient of variation greater than 30% across QC samples, were discarded. Background signals from blank extracted samples were subtracted by default, given the numerous potential sources of endogenous lipid contamination.

#### Bioinformatic Characterization of the LD Lipidome

For the bioinformatic analysis of lipidomic data, data were normalized on total detected lipid amount (mol% of total lipids, all lipid species included), TG species with the same number of carbon atoms and total number of double bonds were pooled together. Tests for differential abundance were performed using an unpaired t-test between replicates. Obtained p-values were then corrected for multiple testing using the method described by Benjamini & Hochberg to obtain FDR-adjusted p-values (*72*). Data were graphically analyzed in R using functions from pcaMethods, pheatmap, and ggplot2 packages and in Prism (Graphpad).

### BRB-sequencing

Total RNA was isolated using RNeasy kit (Qiagen #74134) following the manufacturer’s instructions. The RNA quantity and quality of the RNA was determined using a NanoDrop spectrophotometer (Thermo Fisher Scientific) and the concentration normalized to 100 ng/µl. RNA samples were sent to Alithea Genomics SA (Lausanne, Switzerland) for library preparation and sequencing using highly multiplexed 3′-end bulk RNA barcoding and sequencing (MERCURIUSTM BRB-seq service)(*73*).

#### Library preparation and sequencing

The generation of BRB-seq libraries was performed using the MERCURIUSTM BRB-seq library preparation kit for Illumina and following the manufacturer’s manual (Alithea Genomics, #10813). The library was sequenced on an AVITI from Element Biosciences. Sequencing quality metrics included an average of 7.8 M reads per sample, with 76.1% uniquely mapped reads, 17.9% multi-mapped reads, and 76.36% of reads assigned to annotated genes. Gene detection averaged 14 K genes per sample (range 13.3 K - 14.5 K).

#### Alignment, quantification and data analysis

In this study, we applied STARsolo v2.7.9a to align and quantify the raw barcodes RNA-seq reads against the GRCm38102 (*Mus musculus*) genome (*73*). Utilizing the parameters “—soloUMIdedup NoDedup 1MM_Directional”, and “--quantMode GeneCounts”, we generated raw and UMI-deduplicated count matrices, opting for non-deduplicated counts for subsequent analyses (*74*). Visualization of the differences between conditions was done by transformation of the DESeq2 differential expression data. Regulation is defined by the thresholds log_2_FC = 0.236 and p-value = 0.05.

### RT-qPCR

Cell pellets obtained from cultured NSPCs were immediately snap-frozen on dry ice prior to RNA isolation. Total RNA was extracted using RNeasy® plus mini kit (Qiagen #74134) according to the manufacturer’s instructions. Reverse transcription was performed using the Primescript RT reagent Kit (Takara Bio, RR037A) to generate DNA. qRT-PCR was performed using Power SYBRTM Green PCR Master Mix (#4367659, Fisher scientific) with KiCqStart SYBR green primers for *Cideb* and *beta-actin*, and *Cidea* primer published by Hall et al. (*75*). Relative gene expression levels were calculated using the comparative threshold cycle (ddCT) method, with *beta-actin* used as the endogenous reference gene. Statistical analyses were performed using the calculated ddCt values.

### Oligonucleotides

*SYBR Green RT-PCR primers:*

Cidea_fwd: 5’ ACA GGA GGA CCC GCA CCA AT ’3
Cidea_rev: 5’ GCT GTG CCC TGG TTA CAT GAA C ’3
Cideb_fwd: 5’ CAA GAA CAA CAG AGA AGC AC ’3
Cideb_rev: 5’ CAG TGG ATA CTG ACC TTA G ’3
Actin_fwd: 5’ GAT GTA TGA AGG CTT TGG TC ’3
Actin_rev: 5’ TGT GCA CTT TTA TTG GTC TC ’3

*Oligonucleotides carrying shRNA target sequence (shRNA target sequence in bold):*

shCidea1_fwd: 5’ T**CA GAG TCA CCT TCG ACC TAT A**CT CGA GTA TAG GTC GAA GGT GAC TCT GTT TTT TC ’3
shCidea1_rev: 5’ TCG AGA AAA AA**C AGA GTC ACC TTC GAC CTA TA**C TCG AGT ATA GGT CGA AGG TGA CTC TGA ’3
shCidea2_fwd: 5’ T**AC ACG CAT TTC ATG ATC TT**C TCG AGA AGA TCA TGA AAT GCG TGT TTT TTT C ’3
shCidea2_rev: 5’ TCG AGA AAA AA**A CAC GCA TTT CAT GAT CTT** CTC GAG AAG ATC ATG AAA TGC GTG TA ’3
shCideb1_fwd: 5’ T**GC TAA GGT CAG TAT CCA CTG T**CT CGA GAC AGT GGA TAC TGA CCT TAG CTT TTT TC ’3
shCideb1_rev: 5’ TCG AGA AAA AA**G CTA AGG TCA GTA TCC ACT GT**C TCG AGA CAG TGG ATA CTG ACC TTA GCA ’3
shCideb2_fwd: 5’ T**CC TCT GCA TGG AGT ACC TT**C TCG AGA AGG TAC TCC ATG CAG AGG TTT TTT C ’3
shCideb2_rev: 5’ TCG AGA AAA AA**C CTC TGC ATG GAG TAC CTT** CTC GAG AAG GTA CTC CAT GCA GAG GA ’3
shNT_fwd: 5’ T**CC TAA GGT TAA GTC GCC CT**C TCG AGA GGG CGA CTT AAC CTT AGG TTT TTT C ’3
shNT_rev: 5’ TCG AGA AAA A**AC CTA AGG TTA AGT CGC CCT** CTC GAG AGG GCG ACT TAA CCT TAG GA ’3

### Plasmids

All lentiviral backbone plasmids were bought through Addgene. The pLL3.7 plasmid was a gift from Luk Parijs (Addgene plasmid # 11795; http://n2t.net/addgene:11795; RRID: Addgene_11795) (*76*). The pMD2.G plasmid was a gift from Didier Trono (Addgene plasmid #12259; http://n2t.net/addgene:12259; RRID: Addgene_12259). The plasmid pMDLg/pRRE was a gift from Didier Trono (Addgene plasmid #12251; http://n2t.net/addgene:12251; RRID: Addgene_12251) (*77*). The pRSV-Rev was a gift from Didier Trono (Addgene plasmid #12253; http://n2t.net/addgene:12253; RRID: Addgene_12253) (*77*).

Lentiviral plasmids that express shRNAs were generated by inserting oligonucleotides carrying shRNA target sequence into pLL3.7 vector using HpaI and XhoI restriction sites. Successful integration of each target nucleotide into the vector was confirmed by Sanger sequencing.

### Lentivirus preparation

pLL3.7 shRNA expressing virus was produced as previously described (*78*). Briefly human embryonic kidney 293 T cells were transfected with a pLL3.7, pMD2.G, pMDLg/pRRE and pRSV-Rev using Lipofectamine 2000 (#10696153, Thermo Fisher Scientific) in Opti-MEM (#11520386, Thermo Fisher Scientific). Two days after transfection, the virus was collected by filtering the cell culture medium through a 0.22-μm filter system (#431097, Corning). The filtrate was then concentrated twice using ultracentrifugation at 19 400 rpm. The viral pellet was resuspended in 60 µl of 1 x PBS, aliquoted and stored at -70 °C. Virus titer was estimated by transduction of HEK293T cells with different dilutions of original virus solution and counting the viral colonies three days after transduction.

## Data availability

Proteomics Data are available via ProteomeXchange with identifier PXD078329. BRBseq data will be uploaded to the GEO server and publicly available (GEOxxxx). The lipidomic data are available as lipid concentrations measured across all samples and are provided in table S5.

## Acknowledgements

We thank the Proteomics facility (PAF), the Metabolomics facility (MEP), especially Julijana Ivanisevic and Hector Gallart Ayalla, and the Cellular Imaging Facility (CIF) from the University of Lausanne for their help. We further thank Alithea Genomics for their BRB seq service and analysis of the sequencing data. We also thank Jocelyn Fleurimont for technical help with imaging and data analysis. We further thank our funding sources, the University of Lausanne, and the Swiss National Science foundation (#310030_207587/1 to MK).

## Author contributions

DP and MK developed the concept and planned the study. DP performed all experiments and analyzed and interpreted data. MR and DP developed the LD isolation protocol. DP performed LD isolations and collected samples for omics analysis. MQ performed the proteomics analyses of the LD fraction and helped with experimental design and data analysis/interpretation. DP analyzed and interpreted omics data. DP and MK wrote the manuscript with input from all authors.

---

**Supplementary Figure 1.**
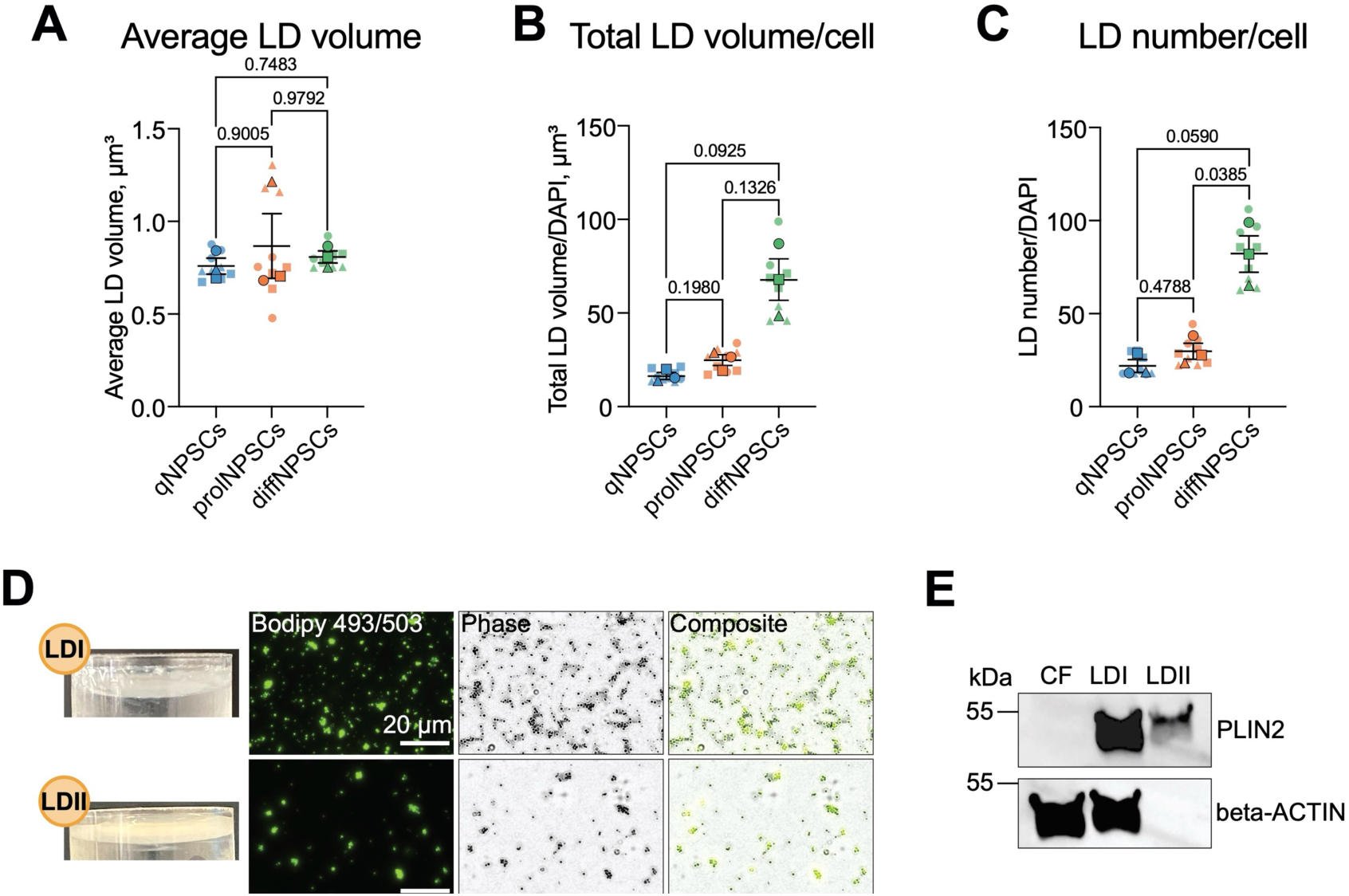
(A-C) Quantification of average lipid droplet (LD) volumes (A), total LD volume/cell (B) and LD number/cell (C) in quiescent neural stem/progenitor cells (qNSPCs), proliferative NSPCs (prolNSPCs), and differentiating NSPCs (diffNSPCs). Each big dot with the same shape represents the average of one independent experiment, each small dot with the same shape represents the average of images coming from the same coverslip. N = 3, n = 9. Mean ± SEM of N = 3, one-way ANOVA. (D) Representative images of the buoyant fractions of LD fraction I and II (LDI and LDII) acquired from LD extraction from prolNSPCs (left). Representative fluorescent and phase contrast images of these fractions stained with BODIPY 493/503 (right). (E) Western blot of cytosolic fraction (CF), LDI and LDII showing amount of PLIN2 and beta-ACTIN proteins detected in these fractions.

**Supplementary Figure 2.**
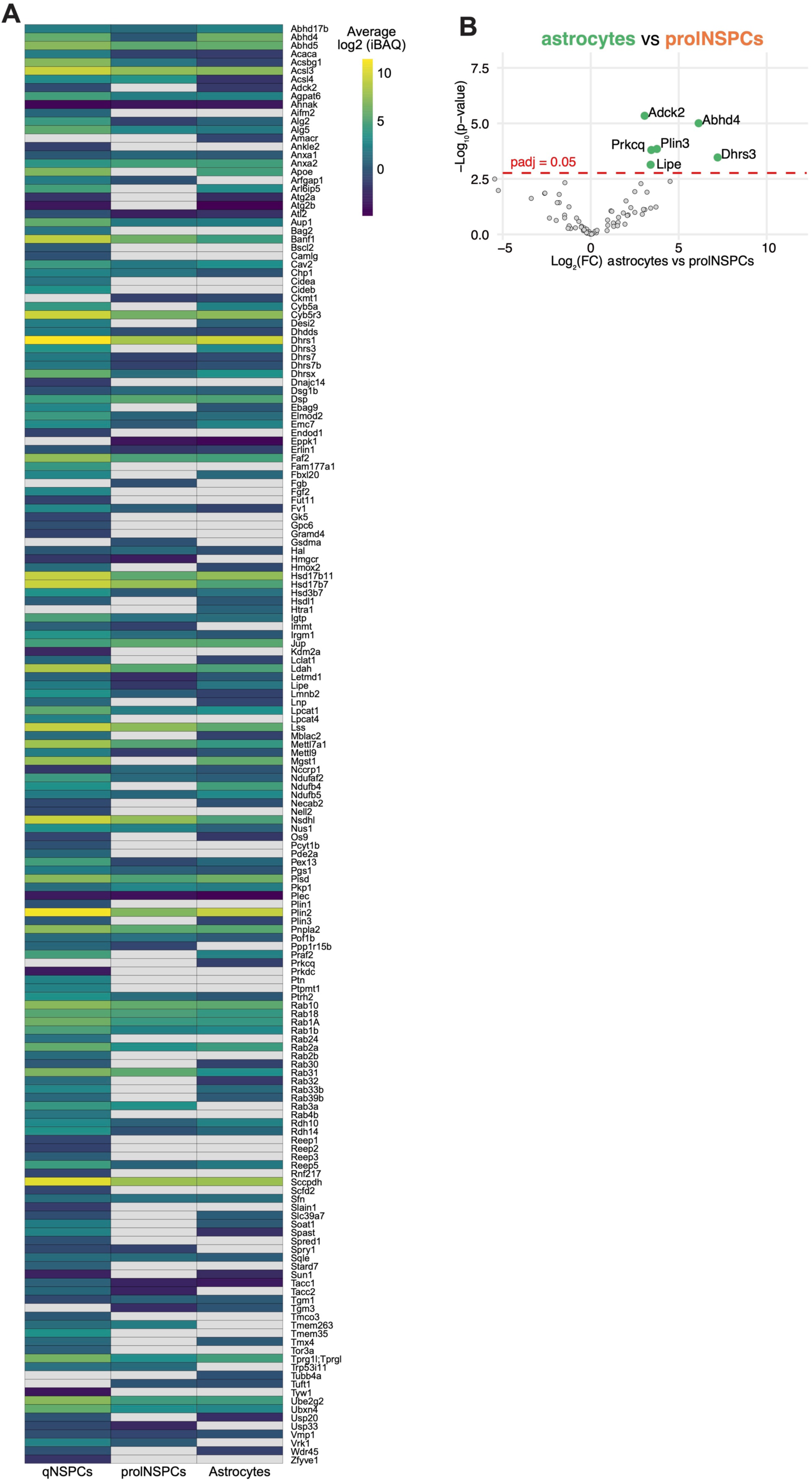
(A) Heatmap of LD protein abundances (average log2iBAQ values), that includes proteins that were at least 2-fold enriched in the LD fraction in at least one cell type. Proteins that were detected in less than 3 out of 4 biological replicates were filtered out and are color-coded gray. (B) Volcano plot based on the high confidence lipid droplet (LD) protein quantification depicting the enrichment of neural stem/progenitor cell (NSPC) LD proteins in prolNSPCs compared to NSPC-derived astrocytes. Significantly upregulated proteins are highlighted in green. Gray represents proteins that did not significantly change in abundance. The p-adjusted (padj) cut-off is 0.05 (adjusted for multiple comparisons). Protein quantification is based on the data collected from four biological replicates.

**Supplementary Figure 3.**
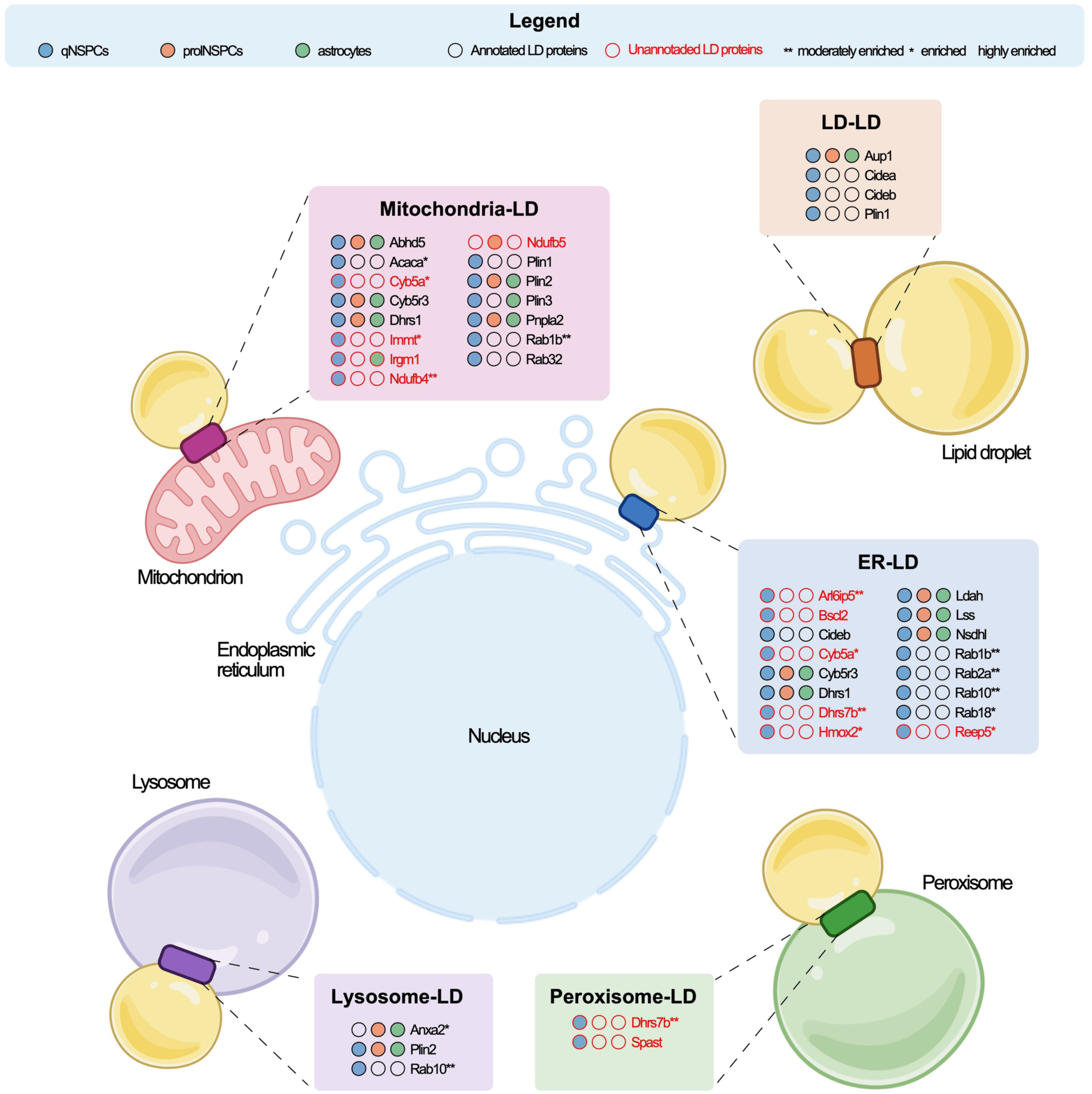
Composite illustration of high-confidence endogenous lipid droplet (LD) proteins identified in quiescent neural stem/progenitor cells (qNSPCs, blue), proliferative NSPCs (prolNSPCs, orange) and NSPC-derived astrocytes (green) that have been described to be involved in the formation of membrane contact sites (MCS). Proteins are color-coded red if they have not been identified as LD proteins in at least one of the datasets used on the LD knowledge portal. Asterisks indicate different cut-off categories: ** for moderately enriched proteins with 1 ≤ log2FC< 2, * for enriched proteins with 2 ≤ log2FC < 3 and no asterisk for highly enriched proteins with 3 ≤ log2FC. ER: endoplasmic reticulum, FC: fold change. Illustration partially created with BioRender.

**Supplementary Figure 4.**
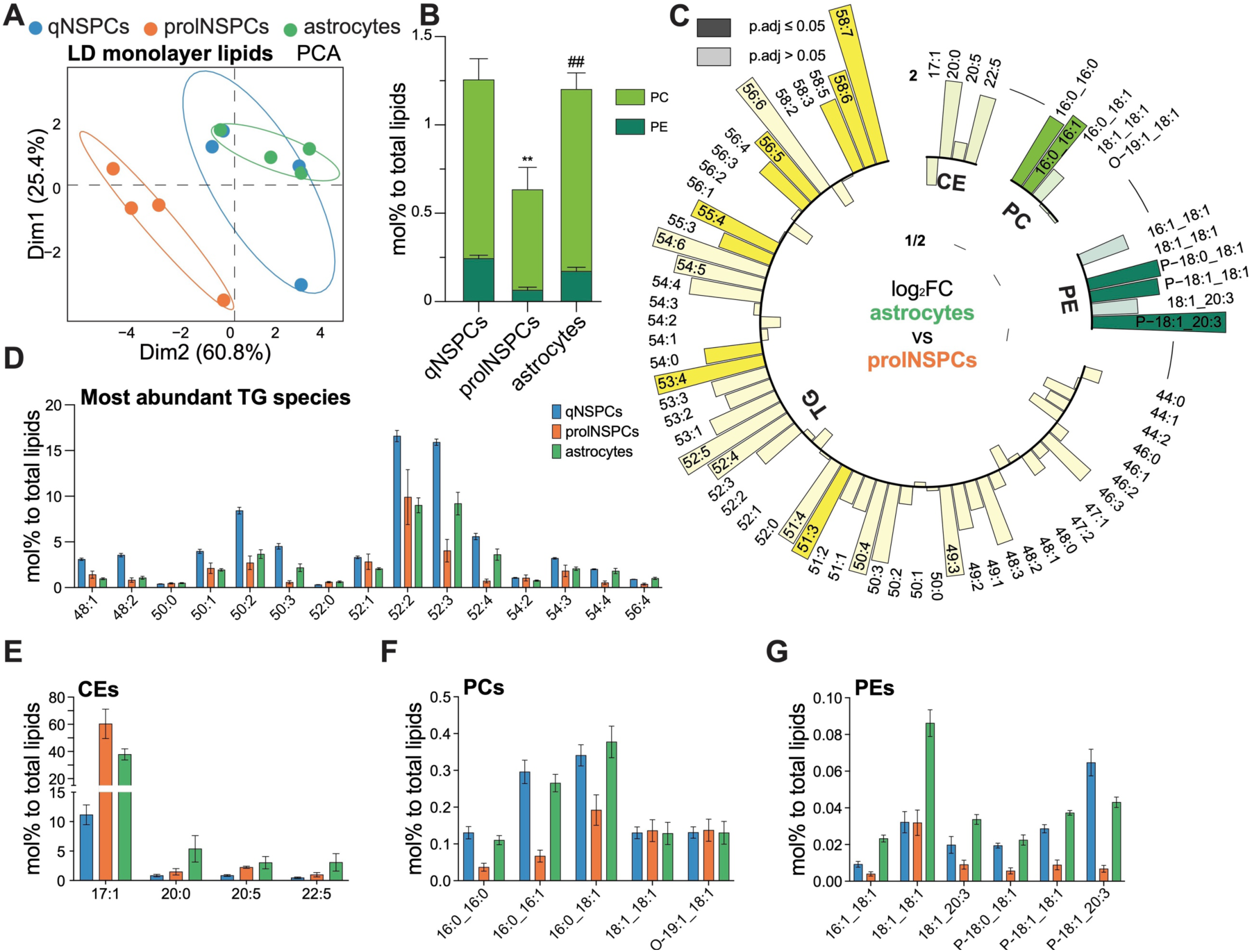
(A) Principal component analysis (PCA) of lipid droplet (LD) samples based on the abundance of identified LD phospholipid monolayer lipids phosphatidylcholines (PC) and phosphatidylethanolamines (PE) in quiescent neural stem/progenitor cells (qNSPCs, blue), proliferative NSPCs (prolNSPCs, orange) and NSPC-derived astrocytes (green). (B) Distribution of LD monolayer lipid species detected in LDs of qNSPCs, prolNSPCs and NSPC-derived astrocytes, color-coded by lipid class. N = 4 samples per condition, mean with SEM, Two-way ANOVA. (C) Circular bar plot based on LD-lipids quantification, depicting logarithmic fold changes (log2FC) of individual lipid species, with their number of carbon atoms : number of double bonds information shown for each lipid NSPC-derived astrocytes compared to prolNSPCs, color-coded by lipid class. Significantly up-(bars directed outside of the circle) or downregulated (bars directed inside the circle) lipids are shown with 100% opacity; bars with 50% opacity represent lipids without significant abundance changes. The p-adjusted cut-off is 0.05 (multiple-comparison adjusted). Lipid quantification is based on four biological replicates.TG = triacylglycerides, CE = cholesterol esters. (D-G) Relative abundances of the fifteen most abundant TG species (D) and all detected CEs (E), PCs (F) and PEs (G) in qNSPCs, prolNSPCs and NSPC-derived astrocytes. N = 4 samples per condition, mean with SEM.

**Supplementary Figure 5.**
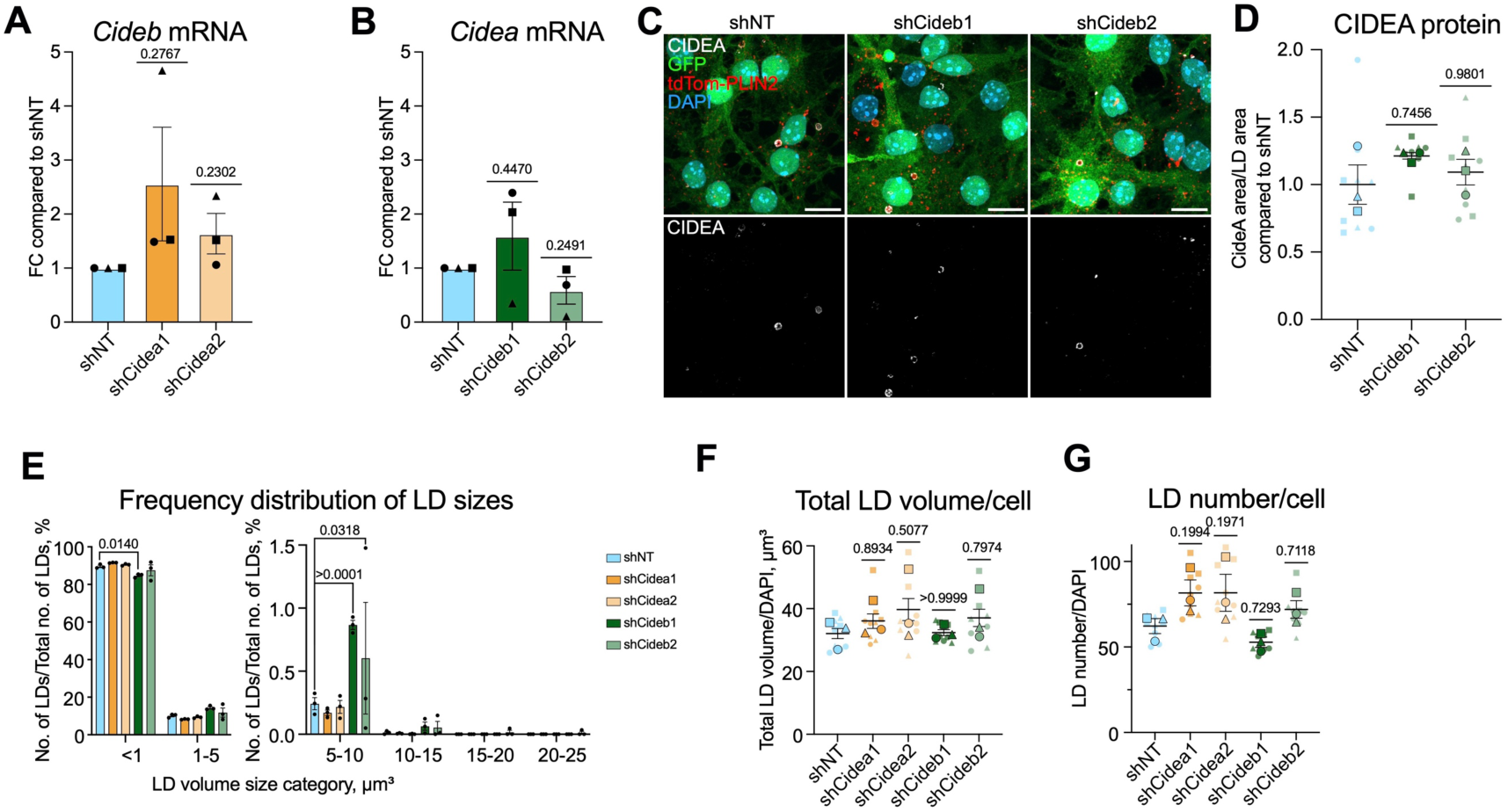
(A, B) Analysis of *Cideb* (A) and *Cidea* (B) mRNA expression by qRT-PCR in quiescent neural stem/progenitor cells (qNSPCs) derived from the tdTom-Plin2+/- mice upon *Cidea* (A) or *Cideb* (B) knockdown (KD) with lentiviral shRNA using two different shRNAs. mRNA levels are normalized on *beta*-*actin* expression. Each dot with the same shape represents an independent experiment, fold change (FC) compared to control non-targeting shRNA (shNT) *Cide* levels ± SEM, one sample t and Wilcoxon test. (C) Representative confocal images of qNSPCs derived from the tdTom-Plin2+/- mice show CIDEA protein levels upon *Cideb* KD using two different shRNAs. Representative images are maximum intensity projections. (CIDEA: CIDEA protein, tdTom-PLIN2: lipid droplets (LDs), GFP: GFP-reporter co-expressed from shRNA-expressing plasmid, DAPI: nuclei). (D) Quantification of CIDEA protein levels upon *Cideb* KD using two different shRNAs. Each big dot with the same shape represents the average of one independent experiment, each small dot with the same shape represents the average of images coming from the same coverslip. N = 3, n = 9. Mean ± SEM of N = 3, one-way ANOVA. (E) Bar graph showing the number of LDs and their respective size frequency distribution upon *Cidea* and *Cideb* KD using two different shRNAs. Each dot represents a separate experiment, with N = 3; n = 9. Two-way ANOVA (factors: condition and LD size distribution) after transformation of data using arcsin(sqrt(Y)) followed by Šídák’s multiple comparisons test. The data representing the mean value ± SEM are depicted for the total number of LDs. (F, G) Quantification of total LD volume/cell (F) and LD number/cell (G) upon *Cidea* and *Cideb* KD using two different shRNAs. Each big dot with the same shape represents the average of one independent experiment, each small dot with the same shape represents the average of images coming from the same coverslip. N = 3, n = 9. Mean ± SEM of N = 3, one-way ANOVA.

**Supplementary Figure 6.**
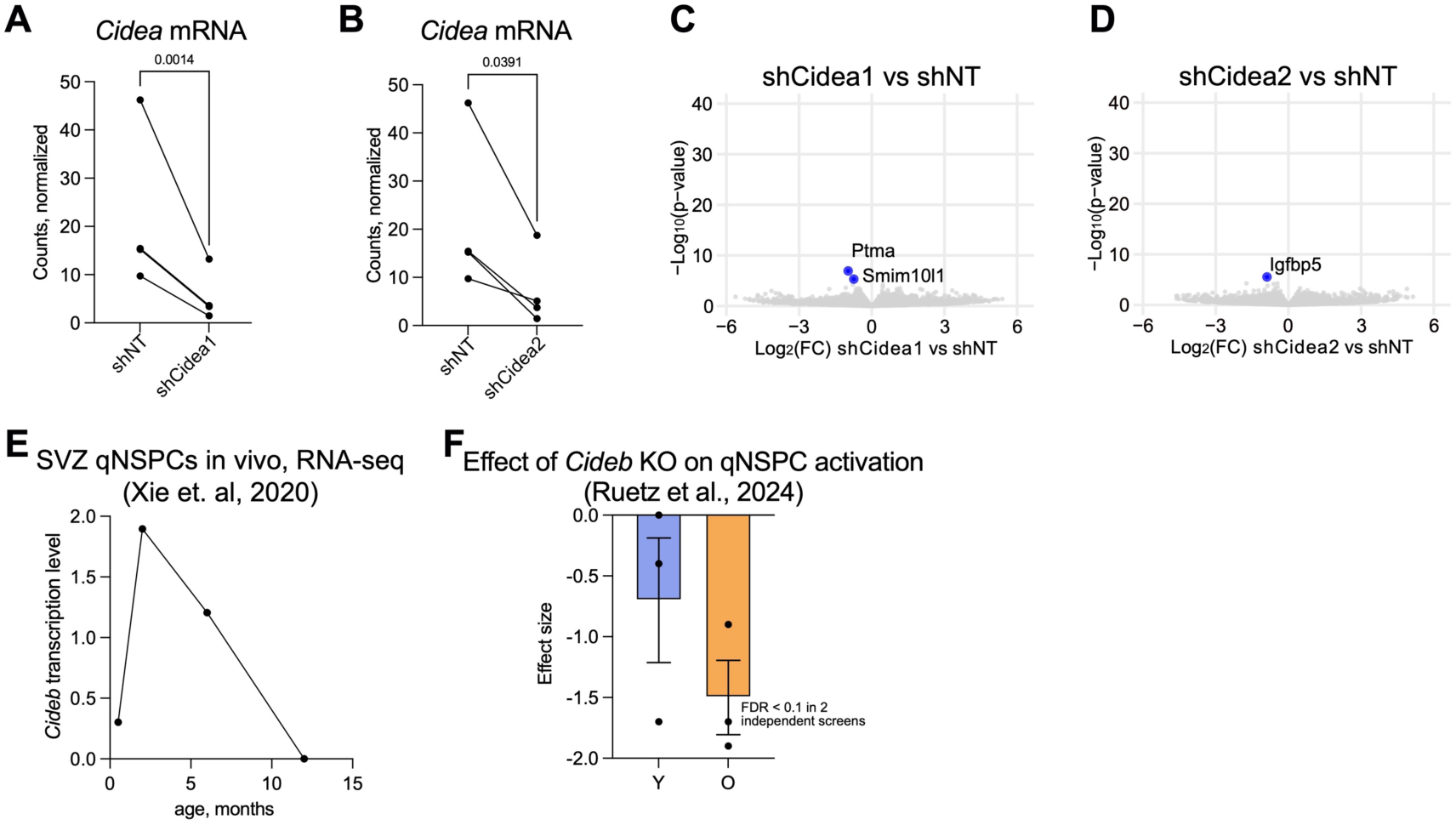
(A, B) Levels of *Cidea* mRNA in quiescent neural stem/progenitor cells (qNSPCs) upon the *Cidea* knockdown (KD) with shCidea1 (A) or shCidea2 (B) shRNAs compared to non-targeting shRNA shNT based on the bulk RNA barcoding and sequencing (BRB-seq) quantification. Each dot represents an independent experiment, whereby the dots belonging to the same experiment are connected by a line. The same shNT control is shown in panels A and B, as both KD experiments were performed in parallel and therefore share a common control. Significance of the KD is tested with a ratio paired t-test. (C, D) Volcano plots based on the BRB-seq quantification depicting the enrichment of genes in qNSPCs upon *Cidea* KD with shCidea1 (C) and shCidea2 (D) compared to shNT. Significantly downregulated genes are shown in blue. Gray dots represent genes that did not significantly change in abundance. The p-adjusted cut-off is 0.05 (adjusted for multiple comparisons). The fold change cut-off is 1.2. Quantification is based on the data collected from four biological replicates. (E) *Cideb* transcriptional levels in qNSPCs isolated from mice brains of different age based on the single-cell RNA-seq data coming from murine subventricular zone (SVZ) NSPCs in vivo collected by Xie and colleagues^45^. (F) Effect size of *Cideb* CRISPR-induced knockout (KO) in qNSPCs extracted from SVZ of young (Y) and old (O) mice on the proliferation rate of NSPCs 14 days after activation. Each dot represents an independent CRISPR screen. KO effect is considered significant if false discovery rate (FDR) is < 0.1 in at least 2 independent screens out of 3. The data is produced by Ruetz and colleagues^60^.

